# Near-Infrared-II Chemo-Optogenetics for Deep-Brain Stimulation

**DOI:** 10.64898/2026.05.13.724823

**Authors:** Hongqiang Yin, Tuanwei Li, Feng Wu, Wuqiao Jiang, Qinhui Yuan, Tao Wang, Yejun Zhang, Chunyan Li, Jiang Jiang, Guangcun Chen, Qiangbin Wang

**Affiliations:** CAS Key Laboratory of Nano-Bio Interface, Jiangsu Key Laboratory of Organoid Engineering and Precision Medicine, Division of Nano-biomedicine and i-Lab, Suzhou Institute of Nano-Tech and Nano-Bionics, Chinese Academy of Sciences, Suzhou 215123, China; School of Nano-Tech and Nano-Bionics, University of Science and Technology of China, Hefei 230026, China; School of Physical Sciences and Technology, ShanghaiTech University, Shanghai 201210, China

**Author notes:** Senior author (Guangcun Chen). Correspondence (G.C.); (Q.W.).

## Abstract

Precise activation of ion channels enables fine-tuned control of neuronal excitability, providing a powerful strategy for dissecting and modulating neural circuits. Although optogenetics offers high spatiotemporal neuronal manipulation with cell-type specificity, its reliance on visible light (400-650 nm) limits tissue penetration to millimeter depths, restricting applications in deep-brain stimulation. Here we report a near-infrared-II (NIR-II)-sensitive calcium channel, HaloNeu, created by genetically fusing a circularly permuted HaloTag (cpHaloTag) to the thermo-sensitive transient receptor potential vanilloid 1 (TRPV1) and covalently conjugating NIR-II photothermal nanotransducers (HPN). The HaloNeu enables non-invasive neuromodulation at depths up to 1.0 cm at ultralow laser power (∼60 mW/cm^2^) and up to 5.0 cm under the safe exposure limit (∼1 W/cm^2^) with 1064 nm laser illumination. Remarkably, HaloNeu maintains stable, on-demand neuron-specific modulation for over two months *in vivo*, providing sustained activation of ventral tegmental area (VTA) circuits and effective alleviation of Parkinsonian symptoms in mouse models. These results establish HaloNeu as a robust and versatile platform for cell-type-specific, deep-tissue, and chronic neuromodulation, with broad implications for neuroscience and neurotherapeutics.

## INTRODUCTION

An ideal neuromodulation strategy requires a synergistic integration of cell-type specificity, high spatiotemporal resolution, deep-tissue penetration, non-invasiveness, and chronic stability^1,2^. Existing neuromodulation modalities—including deep brain stimulation (DBS), transcranial magnetic stimulation (TMS), focused ultrasound (FUS), chemogenetics, and optogenetics—have advanced our capability to manipulate neuronal activity, yet each only partially fulfills these benchmarks^3–5^. DBS, TMS, and FUS lack cell-type-specific precision, increasing the risk of off-target effects^6^. Chemogenetics achieves cell-type-specific modulation via engineered receptors but suffers from limited temporal resolution, typically operating on the scale of minutes to hours^7^. Optogenetics offers exceptional precision by targeting light-sensitive ion channels in specific cells using visible light (400-650 nm), however, tissue scattering and absorption at these wavelengths necessitate invasive optical fibers or high-power lasers, compromising safety^8^. Thus, no current modality fully satisfies the combined requirements of cell-type specificity, deep penetration, non-invasive, and longitudinal stability.

To address these limitations, recent efforts have focused on converting high-penetration physical stimuli—such as magnetic fields, ultrasound, or second near-infrared light (NIR-II, 900-1700 nm)—into thermal, mechanical, or electrical cues to activate endogenous ion channels, including thermo-sensitive transient receptor potential vanilloid 1 (TRPV1), mechano-sensitive TRPV4/Piezo1, and voltage-gated sodium (Na_v_) or potassium (K_v_) channel^9–12^. Notably, NIR-II neuromodulation strategies employing upconversion nanomaterial or photothermal/photomechanical transducers enable wireless modulation of neurons in the ventral tegmental area (VTA) at depths up to 4.5 mm in freely moving mice, eliminating the need for implanted fibers^10,13,14^. Nevertheless, key challenges persist. First, inefficient coupling between transducers and ion channels restricts modulation depths of less than 0.5 cm—a severe constraint that hinders both mechanistic investigation in deep-brain circuits and clinical translation^12,15,16^. Second, conventional antibody-conjugated transducers offer limited ion channel targeting specificity and lack cell-type selectivity, as many target channels (e.g., TRPV1, Piezo1) are expressed in both neurons and glia^17,18^. Third, existing transducers are susceptible to rapid clearance by microglia, glymphatic flow, or enzymatic degradation, limiting effective modulation to less than two weeks^19^.

Here, we introduce HaloNeu, a NIR-II chemo-optogenetic calcium channel. In this system, cpHaloTag-fused TRPV1 (HaloTRP) is selectively expressed in target cells while preserving both TRPV1 channel function and cpHaloTag activity. HaloTag chloroalkane-functionalized NIR-II photothermal nanotransducers (HPN) covalently bind to cpHaloTag, ensuring efficient and specific coupling of NIR-II light to channel gating with subsecond temporal kinetics. This design enables neuromodulation at tissue depths up to 1.0 cm with a 1064 nm laser at ultralow power (∼60 mW/cm^2^) and up to 5.0 cm under a safe exposure limit (∼1 W/cm^2^). Notably, the 5.0 cm depth represents an order-of-magnitude improvement over existing NIR-II approaches and surpassing the performance of reported magnetogenetic and sonogenetic strategies^20–22^. Furthermore, HaloNeu achieves cell-type-specific neuronal activation sustained for up to two months in mice. Collectively, these advances establish HaloNeu as a robust, versatile platform for cell-type-specific, non-invasive, and chronic NIR-II chemo-optogenetics with broad applicability in neuroscience and potential for therapeutic translation.

## RESULTS

### Design of a NIR-II-sensitive calcium channel: HaloNeu

Calcium channels are essential regulators of neuronal excitability, synaptic plasticity, and gene expression^23^. To engineer a calcium channel responsive to NIR-II light, we utilized the thermo-sensitive TRPV1 channel as a structural scaffold and coupled it with high-efficiency NIR-II photothermal nanotransducers. Specific and stable conjugation of these photothermal molecules to TRPV1 is essential for conferring NIR-II sensitivity. We employed the HaloTag self-labeling system, which exploits a modified bacterial haloalkane dehalogenase to form covalent bond with chloroalkane ligands via nucleophilic displacement^24,25^, offering a robust method for precise targeting of TRPV1. Structural analyses indicate that TRPV1 assembles as a tetramer to form a functional calcium ion-permeable pore^26^. Notably, gain-of-function mutations in TRPV channels frequently localize to the S4-S5 loop, the S5-S6 pore region, and the TRP domain, whereas the S1-S4 domain exhibits minimal conformational change during activation and is less prone to mutational perturbation^27,28^. Capitalizing on this structural insight, we rationally inserted a circularly permuted dehalogenase (cpHaloTag) between the S2 helix and the adjacent extracellular S1-S2 loop of each TRPV1 monomer (TRP_m_) to generate a HaloTRP monomer (HaloTRP_m_; Figures S1A and S1B, Figure 1A). The resulting HaloTRP tetramer maintained the overall fold and pore architecture of wild-type TRPV1 (Figure S1C, Figure 1B). Fluorescence imaging confirmed the co-expression of TRPV1 and cpHaloTag in the HaloTRP complex, establishing the foundation for a modular chemo-optogenetic platform (Figure S1D).

**Figure 1.**
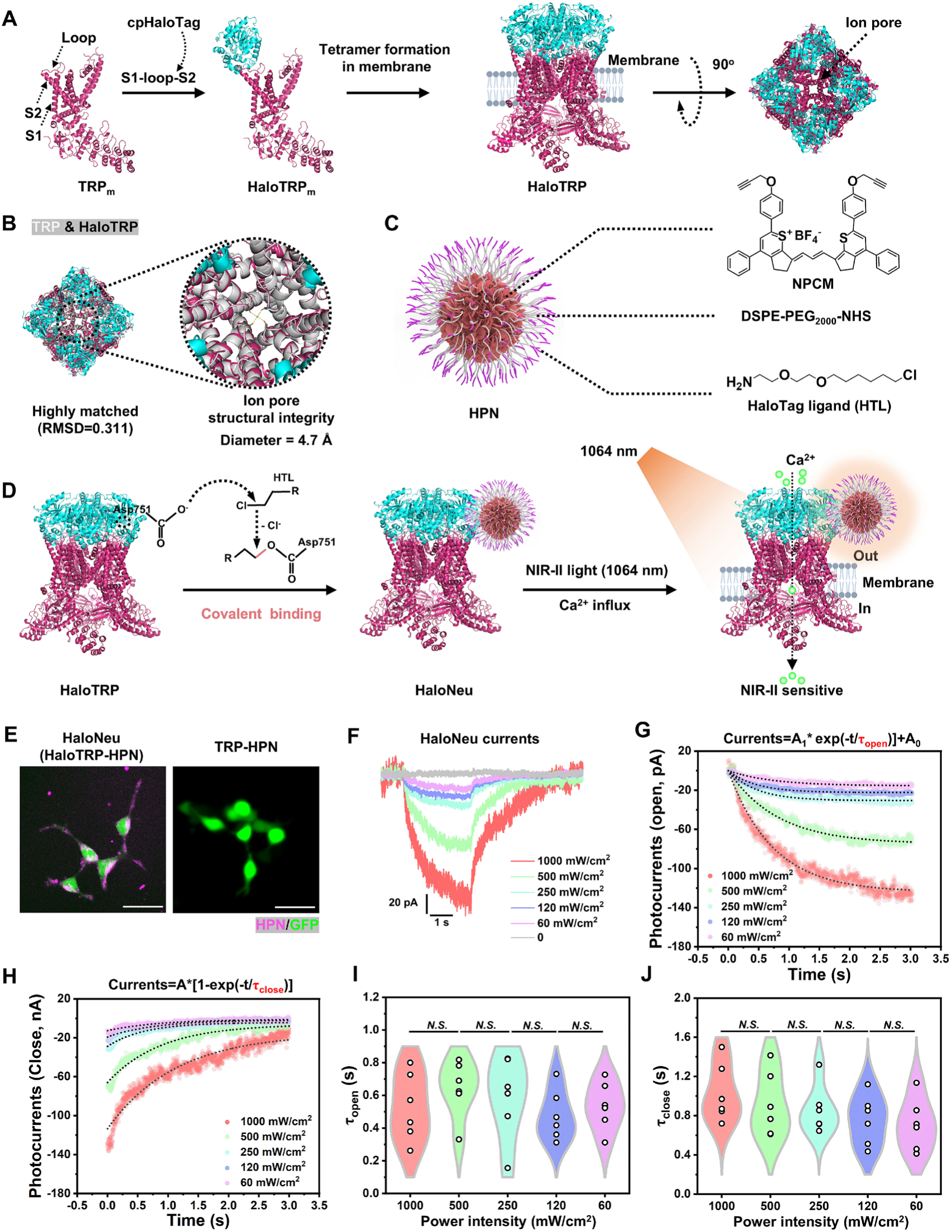
Design and characterization of a biohybrid NIR-II-sensitive calcium channel HaloNeu. **(A)** Schematic of HaloTRP, a fused protein incorporating with TRPV1 and cpHaloTag, where cpHaloTag is inserted between the S1-S2 loop and S2 of TRPV1. **(B)** Structural model of the HaloTRP tetramer, preserving the native TRPV1 fold. **(C)** Composition of HPN, comprising an NPCM core and a surface-functionalized HTL shell. **(D)** Schematic of covalent conjugation between HTL and cpHaloTag in HaloTRP via forming an irreversible ester bond with Asp751, establishing an NIR-II-sensitive calcium channel HaloNeu. **(E)** Specific binding of HPN to HaloTRP-expressing cells. Scale bar, 25 μm. **(F)** HaloNeu currents elicited by 1064 nm light at varying intensities (0-1000 mW/cm^2^). (**G and H**) Mono-exponential fitting of HaloNeu current activation **(G)** and deactivation **(H)**. The time constant τ_open_ and τ_close_ reflect channel opening and closing kinetics. **(I and J)** Analysis of the opening (τ_open_, **I**) and closing (τ_close_, **J**) kinetics of HaloNeu. Data were presented as Mean ± S.D. P ≥ 0.05, no significant difference (*N.S.*).

To render HaloTRP sensitive to NIR-II light, we synthesized HaloTag-ligand (HTL)-functionalized NIR-II HPN. Each HPN incorporates an optimized NIR-II donor-acceptor-donor (D-A-D) photothermal conversion molecular core (NPCM; Figure S2A) and is surface-coated with DSPE-PEG_2000_-HTL to optimize aqueous solubility and biocompatibility (Figures S2B-S2E, Figure 1C). HPN presented a high photothermal conversion efficiency (71% at 1064 nm; Figures S2F-S2I), establishing it as a high-performance nanotransducer ideally suited for optothermal neuromodulation. In addition, HPN possesses fluorescence emission at 555 nm and 1130 nm (Figure S2C), and pronounced NIR-II photoacoustic (PA) signals (Figures S2J and S2K), facilitating multi-modal tracking both *in vitro* and *in vivo*.

To characterize the molecular interface and conjugation efficiency between HPN and HaloTRP, we performed molecular docking simulations and binding specificity assays. The simulations predicted that the chloroalkane ligand engages the conserved catalytic triad (Glu-His-Asp) of HaloTag^29^, forming a covalent ester bond with Asp751 and a hydrogen bond with Gly753 within cpHaloTag (Figure 1D, Figure S3A). Leveraging this mechanism, HPN covalently and stably conjugates to HaloTRP, generating the NIR-II-responsive calcium channel, HaloNeu (Figure 1D). Confocal imaging revealed specific labeling of HaloTRP-expressing cells, with no labeling observed in TRPV1 (TRP) controls, confirming the formation of the biohybrid HaloTRP-HPN complex (Figure 1E). Whole-cell patch-clamp recordings demonstrated that 1064 nm illumination at ultralow power (60 mW/cm^2^) effectively activated HaloNeu, inducing a membrane current increase of 15.22 ± 0.36 pA (Figure 1F), indicative of high NIR-II photosensitivity. Under 1064 nm illumination, the channels exhibited rapid, reversible gating kinetics, with a subsecond opening time constant (τ_open_ = 0.54 ± 0.06 s) and closing time constant (τ_close_ = 0.71 ± 0.10 s) (Figures 1G-1J). Notably, the fast closing kinetics eliminate prolonged after-activation effects often observed in conventional optogenetic ion channels^30^, ensuring precise temporal control.

HaloNeu function as a calcium channel was then evaluated using calcium imaging and electrophysiology. Upon graded 1064 nm illumination, HaloNeu-expressing cells exhibited a marked increase in intracellular calcium compared to TRP-HPN controls (Figures S3B and S3D; Movie S1). Patch-clamp analysis confirmed membrane depolarization driven by rapid photothermal activation (Figures S3C and S3E). Notably, HaloNeu maintained reversible and repeatable gating across multiple laser on/off cycles, with no degradation of electrophysiological properties (Figures S3C).

Collectively, these results establish HaloNeu as a highly sensitive, NIR-II-responsive calcium channel exhibiting subsecond on/off kinetics, reversible gating, and long-term stability, thereby enabling precise and durable chemo-optogenetic neuromodulation.

### HaloNeu stably labels VTA neurons for two months

Following the validation of NIR-II responsiveness *in vitro*, we evaluated the neuronal targeting specificity and longitudinal stability of HaloNeu in the mouse VTA. We stereotaxically delivered an adeno-associated virus (AAV5-hSyn-HaloTRP-3×Flag), encoding neuron-specific HaloTRP, into the VTA. Three weeks later, HPN was injected into the same region. We monitored the integration of the HaloTRP-HPN complex over a 2-month period, simultaneously quantifying the clearance of unbound HPN by microglia (Figure 2A). Immunostaining confirmed robust HaloTRP expression, evidenced by the colocalization of TRPV1 and HaloTag signals within Flag-tagged VTA neurons (Figure 2B). *In vivo* colocalization analysis between HPN and HaloTRP revealed a diffuse HPN signal throughout the VTA at 1 day and 1 week post-injection (Figure 2C). By 2 weeks, however, the diffuse HPN fluorescence was substantially attenuated, leaving a residual signal tightly restricted to Flag-tagged, HaloTRP-positive neurons. The colocalization coefficient reached 0.73 ± 0.020 by 2-week and remained stable throughout the 2-month observation period (0.73 ± 0.013 at 2-month; Figures 2C and 2E). These data demonstrate that HPN undergoes specific, covalent anchoring to HaloTRP, whereas unbound HPN is efficiently cleared, facilitating selective and persistent neuronal labeling. In the control group (TRP-HPN) lacking the cpHaloTag domain, HPN was completely cleared within 2 weeks and showed no colocalization with TRPV1-positive neurons (Figures 2D and 2F).

**Figure 2.**
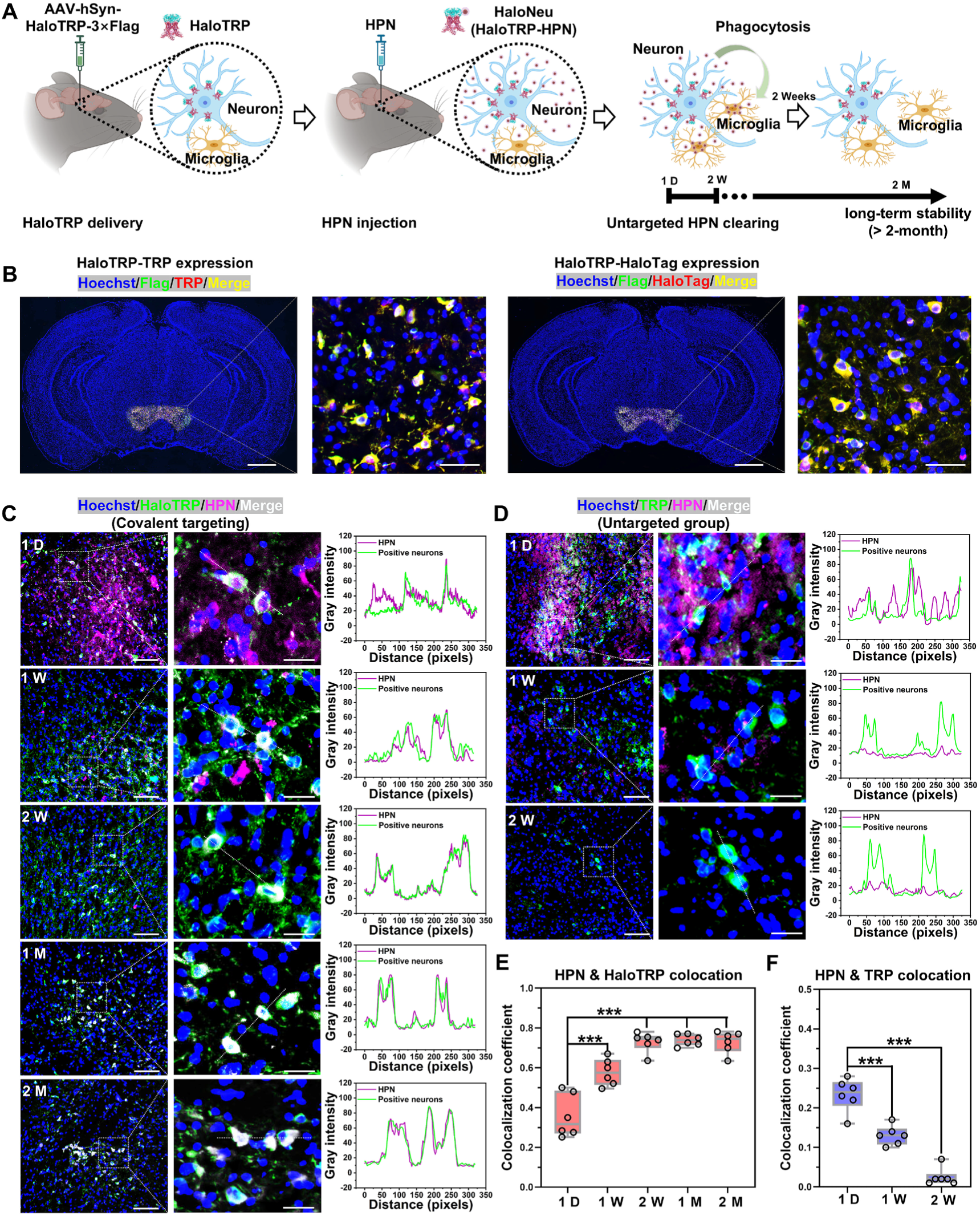
Long-term *in vivo* stability and specificity of HaloNeu. **(A)** Schematic of the experimental workflow: delivery of HaloTRP and HPN, clearance of unbound HPN, and long-term covalent conjugation between HaloTRP and HPN *in vivo*. **(B)** Expression of HaloTRP in VTA region, showing co-expression of TRP and HaloTag proteins. Scale bar (overview), 1 mm. Scale bar (zoom), 30 μm. **(C)** *In vivo* fluorescence imaging showing colocalization of Flag-tagged HaloTRP (green) and HPN (violet, 555 nm emission) over period from 1 day to 2 months. Overlapping fluorescence profiles indicate sustained, specific targeting of HaloTRP by HPN in neurons. Scale bar (overview), 100 μm. Scale bar (zoom), 20 μm. **(D)** *In vivo* fluorescence imaging of Flag-tagged TRP (green) and HPN (violet), demonstrating colocalization from 1 day to 2 weeks. Scale bar (overview), 100 μm. Scale bar (zoom), 20 μm. **(E)** Quantification of the colocalization coefficient between HPN and HaloTRP over time. **(F)** Quantification of the colocalization coefficient of HPN and TRP over time (n=6). Data were presented as Mean ± S.D. *** P < 0.001.

Leveraging the NIR-II photoacoustic (PA) properties of HPN, we performed non-invasive, transcranial PA imaging to track the spatiotemporal distribution of HPN post-injection (Figure S4A). In HaloNeu mice, PA signals within the VTA remained detectable for up to 2 months (Figures S4B and S4D), consistent with immunofluorescence findings. Conversely, both PA and immunofluorescence signals were undetectable after 2 weeks in the TRP-HPN control group, which lacks covalent tethering (Figures S4C and S4E). These results further corroborate that the long-term retention of HPN in HaloNeu mice stems from stable covalent conjugation.

The mechanism underlying the clearance of unbound HPN was further investigated. Immunostaining indicated that microglial phagocytosis is primarily responsive for removing untargeted HPN from the brain (Figures S5A-S5C). In contrast, covalent conjugation anchors HPN to the membrane-bound HaloTRP scaffold, effectively masking them from microglial recognition as "foreign" particles. This stable integration of the HaloTRP-HPN complex (HaloNeu) is therefore essential for achieving precise, sustained neuromodulation while minimizing off-target effects and immune-mediated clearance.

### HaloNeu enables deep-brain and long-term neuromodulation at ultralow power

Following the validation of stable expression and neuronal labeling in the VTA, we evaluated the *in vivo* performance of the HaloNeu. Given that 60 mW/cm^2^ NIR-II (1064 nm) illumination effectively activated HaloNeu *in vitro*, we assessed whether this ultralow power density could modulate neuronal activity in deep-brain regions without inducing thermal damage. NIR-II irradiation (1064 nm, 60 mW/cm^2^, 5 min) induced a minimal, region-confined temperature increase (ΔT = 2.00 ± 0.06 °C) within the VTA (∼4.5 mm depth; Figures 3A and 3B). This thermal elevation was sufficient to activate TRPV1 channels with a latency of ∼3 s (Figures 3C and 3D), consistent with our *in vitro* kinetics and underscoring the high NIR-II sensitivity of the HaloNeu. Importantly, brain regions along the optical path exhibited negligible temperature changes (Figures 3A and 3B), ensuring region-specific and thermally safe neuromodulation.

**Figure 3.**
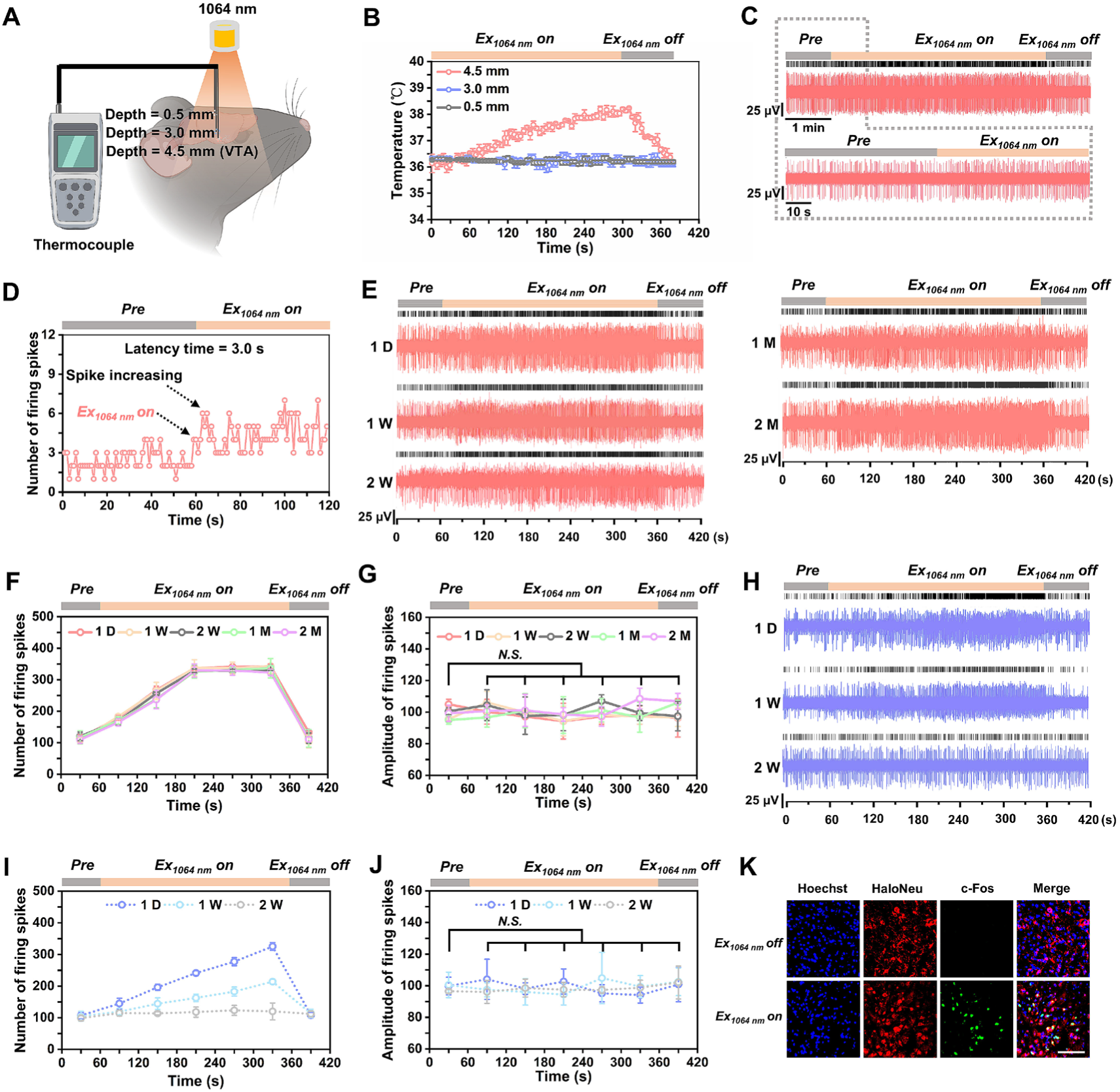
Long-term, stable, neuron-specific, and non-invasive NIR-II neuromodulation *in vivo* using HaloNeu. **(A and B)** Temperature monitoring in VTA and the region across irradiation path under NIR-II neuromodulation (60 mW/cm^2^, n=3). **(C)** Representative electrophysiological traces showing increased firing in HaloNeu neurons upon 1064 nm irradiation (60 mW/cm^2^). **(D)** Quantification of the latency between irradiation onset and increased neuronal spiking in HaloNeu neurons. **(E)** HaloNeu maintained stable long-term neuromodulation efficacy for up to 2 months under ultralow power 1064 nm irradiation (60 mW/cm^2^). **(F and G)** Statistical analysis of neuronal firing spike counts and amplitude before, during, and after 1064 nm illumination across HaloNeu groups (n=3). **(H)** Untargeted control group (TRP-HPN) exhibits only short-term and unstable neuromodulation. **(I and J)** Analysis of spike numbers and amplitude across TRP-HPN groups before, during, and after 1064 nm irradiation (n=3). **(K)** Fluorescence imaging confirming specific c-Fos expression in Flag-tagged HaloNeu neurons 2 months post-delivery, with or without 1064 nm irradiation (60 mW/cm^2^, 5 min). Scale bar, 100 μm. Data were presented as Mean ± S.D.

The HaloNeu strategy contrasts sharply with previous NIR-II photothermal approaches, which typically require approximately 16-fold higher laser powers (1 W/cm^2^) to achieve comparable TRPV1 activation in the VTA. Importantly, the 60 mW/cm^2^ intensity employed here falls within the range routinely applied in near-infrared functional brain imaging, supporting its biocompatibility and suitability for longitudinal or wearable clinical applications. We attribute this high-efficiency, non-invasive deep-brain activation to the covalently formed HaloTRP-HPN complex (HaloNeu), which markedly enhances TRPV1’s NIR-II photosensitivity.

Given the robust *in vivo* retention of HaloNeu and its high optical sensitivity, we examined its long-term functional stability under ultralow-power irradiation. Electrophysiological recordings from 1 day to 2 months post-injection revealed persistent, laser-dependent increases in VTA neuronal firing, and activity levels reverted to baseline upon laser offset, yet the evoked response remained stable throughout the 2-month observation period (Figures 3E and 3F). Furthermore, neuronal spike amplitudes—a reliable proxy for signal propagation integrity^31^—remained unchanged during repeated low-power stimulation, confirming that HaloNeu induces no detectable neuronal damage (Figure 3G). In contrast, neurons in the TRP-HPN control group exhibited transient responses that vanished within 2 weeks, correlating with the microglial clearance of untargeted HPN (Figure 2D, Figures S5C, Figures 3H-3J). No significant activation was observed in neurons expressing TRP alone, HaloTRP alone, or those receiving HPN alone (Figures S6A-S6D). Complementary c-Fos immunostaining—a reliable marker of recent neuronal activity—revealed robust expression in HaloNeu neurons upon 1064 nm stimulation (Figure 3K), corroborating that a stable covalent bond between HaloTRP and HPN is essential for sustained, specific, and low-power neuromodulation.

Collectively, these results establish HaloNeu as a high-performance neuromodulation platform, operating at an ultralow power density (60 mW/cm^2^) with long-term stability (∼2 months) and cell-type specificity, which is far beyond the current non-invasive neuromodulation technologies (Table S1)^32,33^, offering a promising platform for advanced brain research.

### HaloNeu enables remotely, on-demand manipulation mouse behavior

Dopaminergic (DA) neurons, which constitute the predominant neuronal population in the VTA, play a central role in integrating synaptic inputs to encode diverse signals that mediate natural rewards processing, motivation, and pleasure-related behaviors^34^. Based on this framework, we hypothesized that NIR-II irradiation could selectively activate HaloNeu in VTA DA neurons to evoke specific behavioral outputs via DA-dependent neurotransmission (Figure 4A). Immunostaining confirmed predominant expression of Flag-tagged HaloTRP in VTA DA neurons, as evidenced by the high degree of colocalization with tyrosine hydroxylase (TH, Figure 4B). This cell-type-specific targeting provided a foundation for bidirectionally modulating pleasant and excitable behaviors through controlled manipulation of DA neuronal activity.

**Figure 4.**
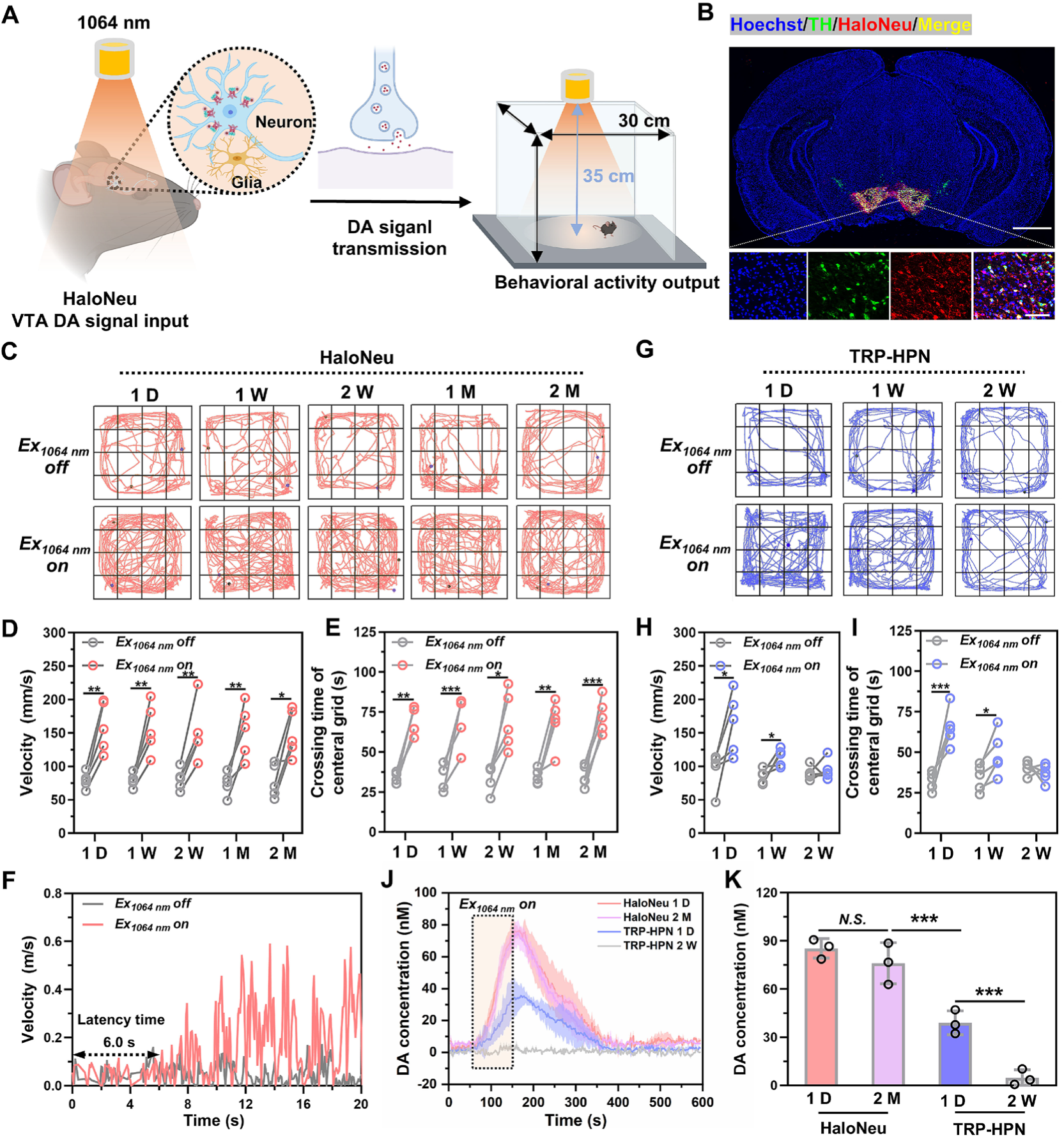
Behavioral modulation of mice using HaloNeu. **(A)** Schematic of HaloNeu-mediated behavioral modulation driven by VTA DA neurotransmission. **(B)** Delivery of HaloNeu (Flag-tagged) into DA neurons (TH-positive) within the VTA region. Scale bar (overview), 1 mm. Scale bar (zoom), 100 μm. **(C)** Representative locomotor trajectories of HaloNeu mice during 1064 nm irradiation (mean power density: 60 mW/cm^2^, beam diameter: 20.6 cm, duration: 5 min). **(D)** Quantification of locomotor velocity in HaloNeu mice under 1064 nm irradiation from 1 day to 2 months post-injection (n=5). **(E)** Time spent in the center zone of the open-field by HaloNeu mice during irradiation across the same period (n=5). **(F)** Latency of behavioral onset following 1064 nm irradiation in HaloNeu mice measured at 2 months post-delivery. **(G)** Locomotor trajectories of TRP-HPN control mice under identical irradiation conditions. **(H and I)** Velocity (**H**) and center zone occupancy (**I**) of TRP-HPN mice under 1064 nm irradiation within 2 weeks post-delivery (n=5). **(J)** DA release monitored by FSCV at VTA synaptic terminals in HaloNeu and TRP-HPN mice under 1064 nm irradiation (60 mW/cm^2^, 90 s) post-delivery (n=3). **(K)** Cumulative DA concentration in VTA synapse after 1064 nm irradiation in HaloNeu and TRP-HPN mice (n=3). Data were presented as Mean ± S.D. *P < 0.05, **P < 0.01, ***P < 0.001.

To evaluate the behavioral consequences of HaloNeu-mediated stimulation, we performed open-field tests in freely moving mice. A 20 W, 1064 nm laser beam was expanded and directed onto the center of the open field arena (30 × 30 × 30 cm^3^), with the beam expander positioned 35 cm above the arena floor. The resulting laser spot diameter was 20.6 cm, yielding an average power density of 60 mW/cm^2^—a level consistent with the parameters used in our *in vivo* electrophysiological recordings (Figure S6E). Under 1064 nm irradiation, HaloNeu mice exhibited a two-fold increase in locomotor velocity (159.42 ± 16.75 mm/s) relative to the non-irradiated condition (79.02 ± 4.05 mm/s), indicating enhanced neuronal excitability driven by VTA DA activation (Figures 4C and 4D; Movie S2). Furthermore, these mice exhibited significantly increased center-arena exploration under NIR-II illumination (Figures 4C and 4E)—a characteristic behavioral signature of DA-mediated reward or anxiolytic responses^35^. Remarkably, this light-evoked modulation persisted for up to 2 months post-injection. Behavioral assessment revealed that even at the two-month time point, the latency to evoke modulation in HaloNeu mice was approximately 6.0 s (Figure 4F)—a timescale comparable to existing NIR-II neuromodulation approaches^9,32^, yet this performance is achieved at an exceptionally ultralow power density of 60 mW/cm^2^—over 16-fold lower than the ∼1 W/cm^2^ typically required by existing NIR-II approaches. This short latency underscores the high temporal precision and rapid responsiveness of the HaloNeu approach in driving behaviorally relevant neural activity. In control experiments, TRP-HPN mice exhibited only transient, unstable behavioral alterations lasting less than 2 weeks (Figures 4G-4I), whereas mice expressing HaloTRP or TRP alone, or receiving HPN only, displayed no significant behavioral changes upon NIR-II exposure (Figures S6F and S6G).

To confirm the dopaminergic basis of these behaviors, we employed fast-scan cyclic voltammetry (FSCV) to measure dopamine release at VTA synaptic terminals following HaloNeu activation (Figures S6H and S6I). In HaloNeu mice, 1064 nm irradiation (60 mW/cm^2^, 90 s) triggered significant DA release, an effect that remained robust for up to 2 months (Figures 4J and 4K). By contrast, TRP-HPN mice exhibited only ephemeral and unstable DA secretion under identical stimulation protocols (Figures 4J and 4K), and no significant DA release was detected in control groups (HaloTRP-, TRP-, or HPN-only; Figure S6J). Together, these results demonstrate that HaloNeu facilitates remote, long-term, and on-demand modulation of reward-related behaviors in freely moving mice through controlled activation of dopaminergic neurons.

### HaloNeu effectively alleviates Parkinsonian motor and psychiatric deficits

Parkinson’s disease (PD) is a progressive, multi-system neurodegenerative disorder characterized by motor impairments—such as tremors and bradykinesia—and non-motor symptoms, including depression^36,37^. The principal pathological hallmark of PD is the degeneration of DA neurons, particularly within the striatum (STR) and the substantia nigra pars compacta (SNc)^38^. Dysfunction within these DA-sensitive circuits is widely considered a key driver of both motor and psychiatric symptoms^39^. In this study, we demonstrate that HaloNeu enables targeted activation of DA neurons, offering a strategy for cell-type-specific, long-term, and stable deep-brain stimulation. Using a PD mouse model, we investigated whether this remote NIR-II optogenetic approach could serve as a viable alternative to conventional DBS—which requires invasive electrode implantation^40^—for intervening in multiple PD-related pathological symptoms.

Given the established role of VTA DA-associated circuits in PD pathology, we examined whether the HaloNeu approach could selectively engage pathways connecting the VTA to the M2 cortex and the nucleus accumbens (NAc) shell to rectify tremor and bradykinesia^41,42^ (Figure 5A). In 1-methyl-4-phenyl-1,2,3,6-tetrahydropyridine (MPTP)-induced PD mice^43^, we observed DA neuron loss in the STR and SNc (Figure S7A), alongside established tremor activity (electromyography monitoring; Figure S7B), bradykinesia, and depression-like behaviors (open field test; Figures S7C and S7D). To elucidate whether the HaloNeu approach can specifically modulate these specific circuits, we conducted whole-brain c-Fos immunostaining. Results confirmed that 1064 nm irradiation successfully activated VTA HaloNeu-expressing (Flag-tagged) neurons in PD mice (Figure 5B), triggering widespread neuronal activation in downstream targets, including the M2 cortex and the NAc shell (Figure 5C). These findings demonstrate that the HaloNeu approach precisely engages VTA-associated neural networks implicated in PD pathophysiology.

**Figure 5.**
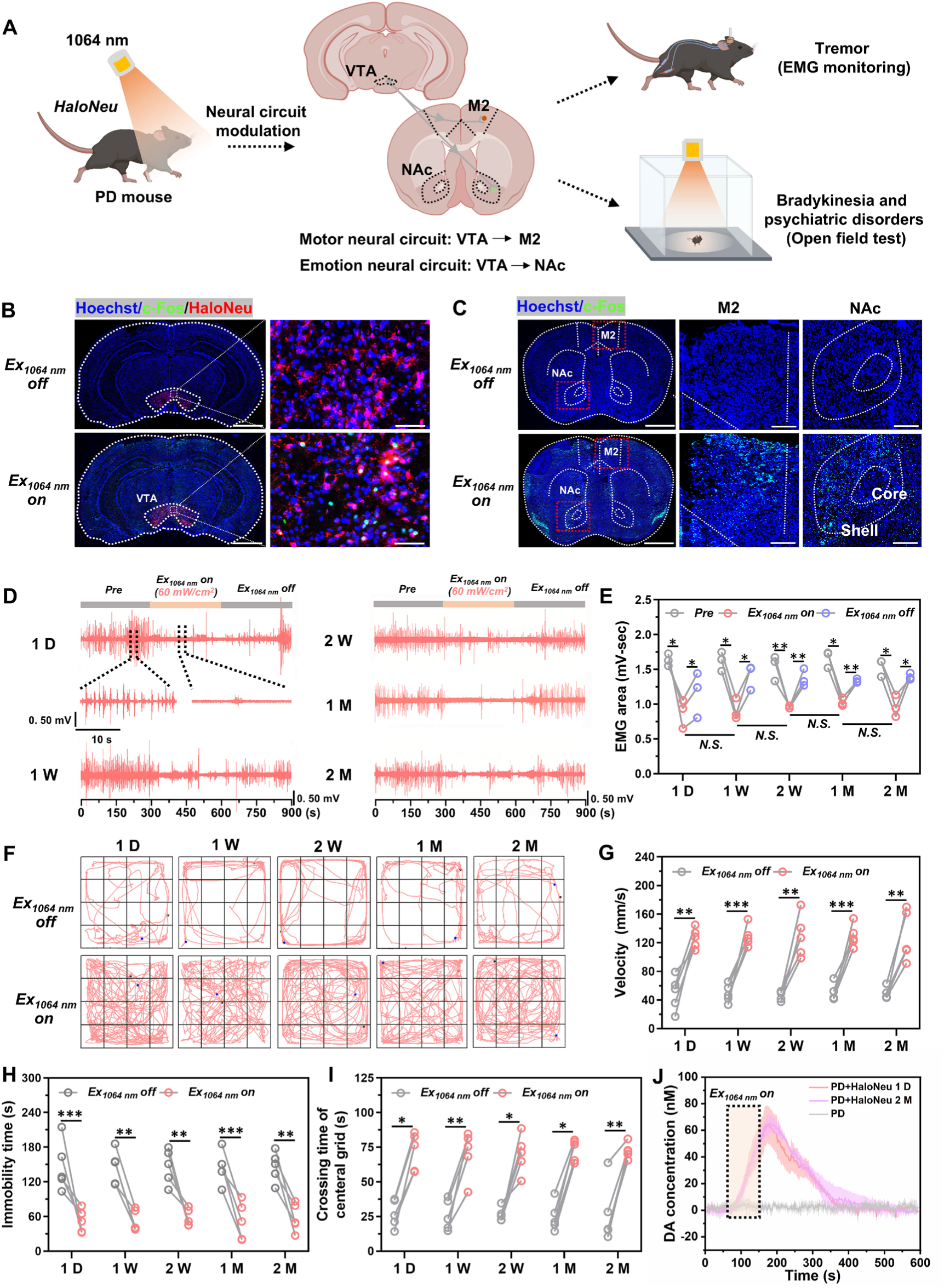
The HaloNeu approach ameliorates motor dysfunction and psychiatric disorders in PD mice via VTA DA-triggered neural circuits activation. **(A)** Schematic illustrating the mechanism by which HaloNeu ameliorates symptoms of PD in mice through activation of VTA DA-triggered neural circuits. **(B)** Selective activation of VTA HaloNeu neurons (Flag-tagged) in PD mice upon 1064 nm irradiation (60 mW/cm^2^, 5 min). Scale bar (overview), 1 mm. Scale bar (zoom), 100 μm. **(C)** Robust c-Fos expression in M2 cortical and NAc shell regions, induced by VTA neuronal activation via the HaloNeu approach in PD mice. Scale bar (overview), 1 mm. Scale bar (zoom), 100 μm. **(D)** EMG recording of tremor activity in PD mice following VTA delivery of HaloNeu, under 1064 nm laser irradiation (60 mW/cm^2^ at scalp, 5 min). **(E)** Quantification of EMG area across groups (n=3). **(F)** Behavioral trajectories of PD HaloNeu mice during 1064 nm irradiation over a 2-month period (mean power density: 60 mW/cm^2^, irradiation diameter: 20.6 cm, exposure time: 5 min). **(G-I**) Statistical evaluation of locomotor velocity (**G**), immobility duration (**H**), and center grid occupancy (**I**) in PD HaloNeu mice under 1064 nm irradiation from 1 day to 2 months post-treatment (n=5). **(J)** FSCV measurement of DA release at the VTA DA neuron synapse in PD HaloNeu mice, assessed 2 months post HaloNeu delivery under NIR-II neuromodulation (n=3). Data were presented as Mean ± S.D. *P < 0.05, **P < 0.01, ***P < 0.001.

To determine the efficacy of the HaloNeu approach in mitigating tremors—a hallmark motor symptom of PD—we performed EMG analysis (Figure S7E). In PD mice with HaloNeu delivered to the VTA, 1064 nm irradiation (60 mW/cm^2^) normalized aberrant firing patterns, with tremor activity consistently re-emerging upon laser cessation (Figure 5D, Movie S3). Notably, the HaloNeu approach enabled stable, real-time tremor intervention throughout the 2-month observation period (Figures 5D and 5E). We further evaluated the effect of the HaloNeu approach on motor activity and depression-like behaviors in PD mice using an open field test. Under 1064 nm irradiation, PD mice exhibited significantly increased velocity, indicating a marked alleviation of bradykinesia (Figures 5F and 5G; Movie S4). Furthermore, immobility time was also reduced under irradiation, consistent with effective intervention in both depression and bradykinesia (Figure 5H). PD mice spent more time in the central area of the open field, suggesting that neuromodulation conferred anti-depressive effects and enhanced exploratory behavior (Figure 5I). These therapeutic effects were maintained for 2 months, enabling on-demand management of both motor and psychiatric symptoms of PD (Figures 5F-5I). To probe the underlying mechanism, we measured DA release in the NAc region of PD mice using FSCV. DA release increased progressively under 1064 nm stimulation, with stable potential of DA secretion over time (Figure 5J). These results indicate that the HaloNeu approach provides a long-term, stable intervention for both motor and psychiatric deficits associated with PD.

### HaloNeu enables centimeter-scale, deep-tissue neuromodulation

Our results established that wide-field NIR-II irradiation (1064 nm, 60 mW/cm^2^) effectively activates VTA neurons located ∼4.5 mm beneath the mouse skull, far exceeding previously reported approaches that required substantially higher laser irradiation densities^9^. To delineate the capability of HaloNeu for deep brain stimulation, we developed an optical phantom using stacked fish tissue to simulate deep-brain light propagation. Systematic characterization of 1064 nm transmittance confirmed that fish tissue closely recapitulates the attenuation profile of the mammalian brain, with nearly equivalent transmittances of 34.04% and 31.11%, respectively (Figures 6A and 6B), indicating comparable effective photon attenuation characteristics, thereby supporting the use of fish tissue as a physiologically relevant optical surrogate for mammalian brain tissue in penetration studies. Linear regression analysis further demonstrated that power density scales predictably with incident intensity in this model, confirming its reliability for deep-tissue modeling (Figure 6A). Leveraging this validated phantom, we demonstrated that even at a minimal irradiance of 60 mW/cm^2^, HaloNeu reliably evokes robust VTA neuronal firing through a total tissue thickness of 1.0 cm (including 6 mm of stacked phantom; Figures 6D-6F).

**Figure 6.**
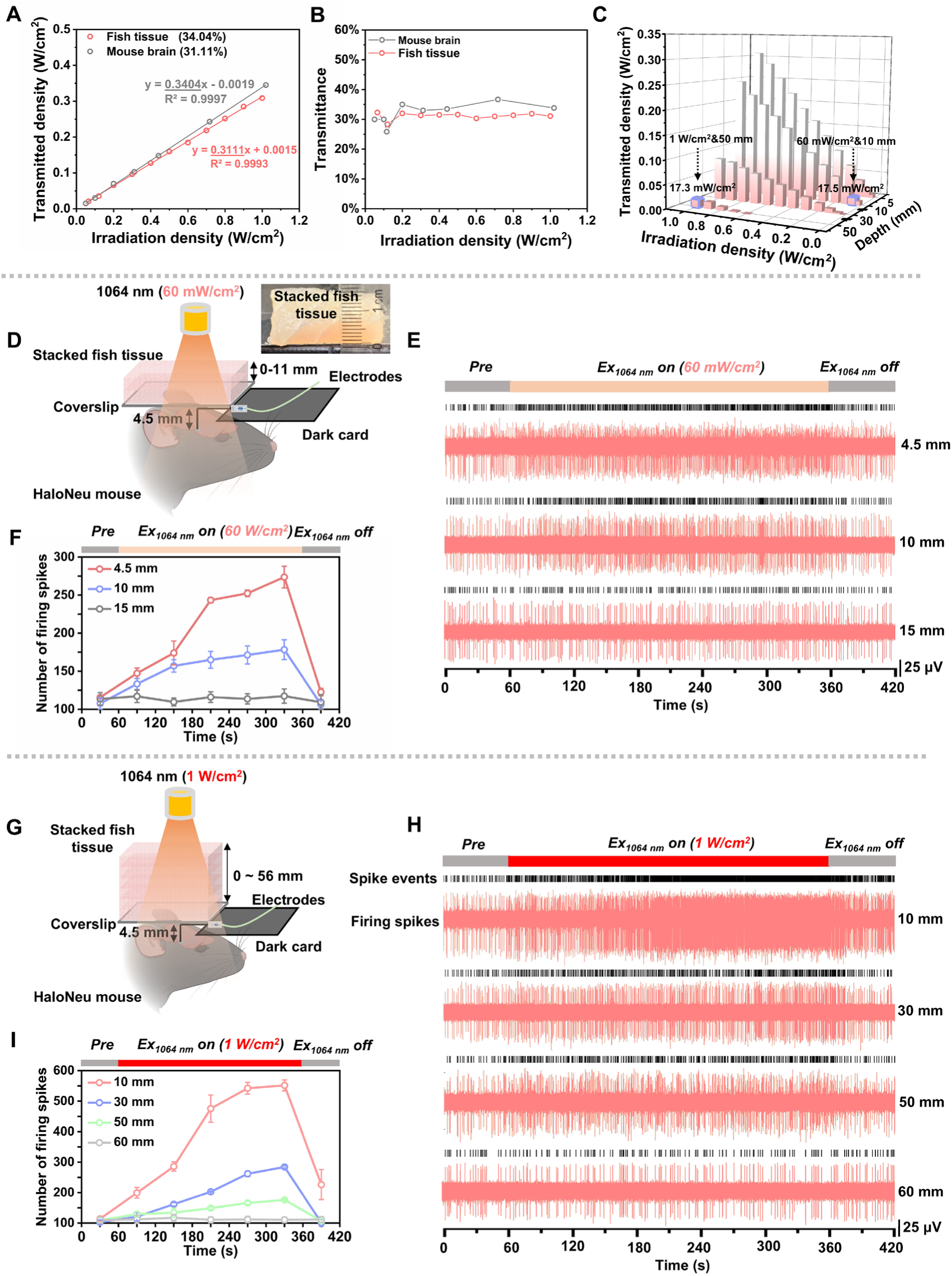
HaloNeu achieves brain stimulation at centimeter-scale depths. **(A and B)** Transmitted power density and transmittance of 1064 nm light at varying incident power density across mouse brain and fish tissues, respectively (n=3). **(C)** Assessment of 1064 nm light penetration through tissue. **(D)** Schematic for evaluating the efficacy of NIR-II neuromodulation through tissue-mimicking layers of varying thicknesses at 60 mW/cm^2^. **(E)** HaloNeu mediates non-invasive neuromodulation at centimeter-scale depths using ultralow power of 1064 nm (60 mW/cm^2^). **(F)** Statistical analysis of firing spike numbers at varying tissue depths (n=3). **(G)** Schematic for evaluating NIR-II neuromodulation efficacy at varying tissue thickness using a fish tissue-mimicking strategy (1.0 W/cm^2^). **(H)** The HaloNeu approach supports unprecedented deep-tissue neuromodulation, enabling neuronal activation at depths up to 5.0 cm. **(I)** Quantification of neuronal firing spike counts across tissue depths under 1064 nm irradiation (1.0 W/cm^2^, 5 min, n=3). Statistical data were presented as Mean ± S.D.

To assess the translational scalability for larger species or clinical intervention, we next evaluated the platform’s performance at an incident intensity of 1 W/cm^2^, which remains strictly within the safety thresholds defined by the International Commission on Non-Ionizing Radiation Protection (ICNIRP)^44^ and has been widely used in recent studies on neuromodulation and photothermal therapy^14,45–47^. Quantitative characterization of depth-dependent attenuation kinetics (Figure 6C) revealed a critical scaling principle for NIR-II light propagation: the residual power density delivered through 5.0 cm of tissue at 1 W/cm^2^ (∼17.3 mW/cm^2^) maintains a reciprocal parity with the intensity delivered through 1.0 cm at 60 mW/cm^2^ (∼17.5 mW/cm^2^). Consistent with this physical parity, HaloNeu facilitated robust and repeatable neuronal activation at unprecedented tissue depths of up to 5.0 cm (including 4.5 cm of stacked phantom; Figures 6G-6I). Notably, this 5.0 cm operational range represents a ten-fold improvement over existing NIR-II platforms (≤ 4.5 mm) and significantly exceeds the reported penetration limits of current magnetogenetic (∼4.5–4.75 mm) and sonogenetic (∼4.5 mm) technologies (Table S1)^14,20,21,48^. This supra-modulatory margin underscores the potential of HaloNeu for large-animal studies and clinical translation.

## DISCUSSION

In this study, we engineered a NIR-II-sensitive calcium channel, HaloNeu, by covalently coupling a genetically encoded HaloTRP channel (TRPV1 fused to cpHaloTag) with NIR-II HPN, thereby establishing a noninvasive chem-optogenetic neuromodulation strategy. The HaloNeu approach achieves centimeter-scale neuromodulation (1 cm at 60 mW/cm^2^; 5 cm at 1 W/cm^2^), cell-type specificity, and rapid temporal kinetics (∼0.5 s channel opening, ∼3 s neuronal activation, ∼6 s behavioral modulation) under 1064 nm irradiation at ultra-low power densities (60 mW/cm^2^), while maintaining robust *in vivo* stability for over two months. Collectively, these advancements surmount the critical barriers of depth, safety, and longitudinal stability, positioning HaloNeu as a transformative interface for systems neuroscience and future clinical neurotherapeutics.

### Mechanistic basis for centimeter-scale penetration of the HaloNeu approach

Limited tissue penetration remains a fundamental barrier to the clinical translation of optogenetics^49,50^. Conventional visible-light optogenetics typically achieves <1 mm penetration^51^, and existing NIR-II neuromodulation approaches generally remain confined to <5 mm^10,13,14,32^ (Table S1). HaloNeu overcomes this limitation by enabling unprecedented centimeter-scale neuromodulation (1 cm at 60 mW/cm^2^; 5 cm at 1 W/cm^2^), representing approximately an order-of-magnitude improvement over current NIR-II strategies based on antibody-targeted ion channels or analogous approaches.

We attribute this substantial enhancement in tissue penetration to the covalent anchoring of HPN to TRPV1 via the HaloTag self-labeling system, which optimizes both stimulus specificity and thermal responsiveness. First, unlike antibody-based targeting strategies that may sterically hinder or unpredictably perturb functional gating domains, our rational insertion of the cpHaloTag between the S2 helix and the adjacent extracellular S1-S2 loop preserves the native pore architecture and intrinsic thermo-sensitivity of the TRPV1 scaffold (Figure 1A and 1B; Figure S1C). By spatially segregating the targeting module from the functional core, this architecture minimizes structural interference and ensures optimal heat-induced gating.

Second, HaloNeu exhibits high energy-transfer efficiency. The covalent linkage between HPN and HaloTRP minimizes the thermal diffusion path in nanoscale and restricts spatial heat dispersion, defined by the PEG-2000 linker (∼4.0 nm in aqueous solution)^52^. To elucidate this ultrafast thermal transport, we employed the Guyer-Krumhansl (C-K) model, which accurately describes confined, non-Fourier heat propagation in nanoscale biological systems^53,54^. The C-K model is as follows:

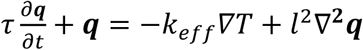

Where *τ* represents the thermal relaxation time, describing delayed heat flux response to temperature gradients; *t* denotes time, representing the progression of the heat transfer process; *k*_*eff*_ represents the effective thermal conductivity of the surrounding media; *l* is the transport length scale; ***q*** is the heat flux; *V̅T* is the temperature gradient.

In HaloNeu system, the covalent linkage between HPN and HaloTRP defines an ultrashort transport length (*l* ≈ 4 nm, corresponding to the PEG-2000 linker), meaning that heat only needs to travel a few nanometers. Scaling analysis of the G-K equation shows that the relaxation time *τ* is comparable to the characteristic heat propagation time across the transport length *l*. In this nanoscale regime, this leads to *τ* ≈ *l*/*v*, where *v* is the effective propagation velocity of thermal energy carriers (∼10^3^ m/s in water), yielding a picosecond-scale time for heat transfer. This picosecond-scale thermal transport occurs on a timescale much faster than the biological response, allowing the system to be treated in the long-time limit (*t* ≫ *τ*), where the G-K model reduces to its Fourier-like steady-state form. Under this condition, the heat flux can be expressed as *q* ≈ −*k*_*eff*_∇*T*, from which an order-of-magnitude estimate of the temperature variation can be obtained as Δ*T*∼*q* · *l*/*k*_*eff*_, where *k*_*eff*_ is used the thermal conductivity of water 0.6 W·m^-1^·K^-1^. This analysis indicates the resulting temperature variation across the molecular coupling distance is on the order of 10^-5^-10^-4^ °C.

This picosecond thermal coupling enables rapid activation of HaloTRP by HPN-generated heat, resulting in subsecond channel kinetics of HaloNeu with an opening time constant (τ_open_ = 0.54 ± 0.06 s) and a closing time constant (τ_close_ = 0.71 ± 0.10 s). Moreover, the minimal thermal attenuation (10^-5^-10^-4^ ℃) further ensures that the energy required to activate HaloNeu is kept to a minimum, which is particularly ciritical for avoiding undesired effects or thermal damage in biological systems. Collectively, these findings elucidate the nanoscale-confined, ultrafast, and low-loss thermal transport mechanism of the HaloNeu system, providing a direct kinetic basis for its ultralow-power activation, high photosensitivity, and feasibility for centimeter-scale neural modulation, while establishing a strong link between the thermal transport characteristics and electrophysiological phenotypes of the composite channel.

Third, the high specificity of the covalent interaction between HaloTRP and HPN mitigates off-target activation. Endogenous thermosensitive channels—such as inhibitory potassium channels (e.g., TREK-1)^55,56^—and surrounding glial populations are often susceptible to non-specific heating^57^, which can counteract neuronal depolarization or introducing network-level modulation that diminishes activation efficiency. By restricting photothermal activation exclusively to HaloTRP-expressing neurons, HaloNeu minimizes these detrimental off-target effects and maximizes the efficiency of neuronal depolarization.

### Therapeutic potential of HaloNeu for neurological disorders

The synergy of cell-type specificity, centimeter-scale penetration, and rapid temporal kinetics positions HaloNeu as a transformative paradigm for optogenetic intervention in neurological diseases. As proof-of-concept, we demonstrated that the HaloNeu approach enables precise control of projection-defined neural circuits originating from the VTA, including the VTA-to-NAc and VTA-to-M2 cortex pathways. Targeted modulation of these circuits leads to rapid tremor suppression and significant reversal of bradykinesia and depression-like behaviors in Parkinson’s mice. These therapeutic outcomes arise from circuit-specific activation of VTA projections, which maximizes clinical efficacy across both motor and psychiatric domains while circumventing the off-target effects typically associated with systemic or non-targeted stimulation.

Crucially, the 5 cm penetration depth achieved by HaloNeu is sufficient to access deep brain regions without the requirement for implanted optical fibers^58^, thereby eliminating risks associated with surgical intervention, such as tissue trauma and chronic neuroinflammation^59,60^. This noninvasive, cell-type-specific, transcranial neuromodulation strategy offers a robust framework for regulating complex cortical and subcortical circuits governing cognition, motor control, and affective states in higher-order species. Furthermore, HaloNeu maintains 1 cm penetration at an irradiance of 60 mW/cm^2^—well within the safety thresholds for chronic biological exposure (≤100 mW/cm^2^)^61^—is highly significant. Given that peripheral targets, such as the vagus nerve, typically reside within 1 cm of the skin surface^62^, HaloNeu facilitates the development of long-term, wearable, non-implantable neuromodulation platforms for disorders such as depression and epilepsy^63^, substantially broadening its translational scope beyond central nervous system applications.

### An extensible and multimodal neuromodulation platform

Beyond the specific TRPV1-based implementation, this work establishes the HaloTag as a modular scaffold for advanced neuromodulation. The fusion of cpHaloTag to TRPV1 generates HaloTRP, a biohybrid channel that can be functionalized with HPN to confer NIR-II sensitivity. More broadly, the covalent interaction between the HaloTag and its ligand (HTL) offers a robust anchoring strategy for a variety of nanotransducers, enabling the rational design of genetically encoded ion channels responsive to diverse physical stimuli—including magnetic, ultrasonic, and optical cues. This HaloTag-based framework is inherently generalizable. By fusing the cpHaloTag to different ion channel families, including both excitatory and inhibitory channels, or to G protein-coupled receptors^64,65^, this platform supports multifunctional and combinatorial neuromodulation. Such modularity enables coordinated, multi-channel, and multi-region control with high spatiotemporal precision and chronic stability, providing a versatile toolkit for next-generation biohybrid neural interfaces.

### Limitations of the study

While HaloNeu represents a substantial advancement in non-invasive neuromodulation, several factors necessitate further consideration. Although we demonstrated centimeter-scale penetration in tissue phantoms and robust functional outcomes in murine models, the efficacy, safety profile, and precision of this platform in large-animal brains require systematic validation. Owing to species-specific differences in endogenous TRPV1 sequences, viral transduction efficiency, and brain scaling, a dedicated large-animal study is required and is currently underway. Furthermore, the current requirement for exogenous HPN replenishment to maintain modulation beyond two months may constrain chronic therapeutic applications. Further development of blood-brain barrier-permeable photothermal transducers, which would facilitate minimally invasive re-labeling of HaloTRP-expressing neurons, is essential to enhance the platform’s durability and clinical scalability. Finally, while our therapeutic assessment focused on motor and depression-related deficits in Parkinson’s models, the broader applicability of HaloNeu in addressing cognitive dysfunction, sleep disturbances^66,67^, and other complex neurological comorbidities remains to be developed.

## RESOURCE AVAILABILITY

### Lead contact

Requests for further information and resources should be directed to and will be fulfilled by the lead contact, Qiangbin Wang.

### Materials availability

Plasmids generated in this study have been deposited to Addgene (HaloTRP, ID: 231245). Moreover, Other materials in our study are also available from the lead contact upon request.

### Data and code availability

The original datasets are accessible at Mendeley under http://doi.org/10.17632/sy6drj5cfp.1. This study does not report original code. Any additional information required to reanalyze the data reported in this paper is available from the lead contact upon request.

## ACKNOWLEDGMENTS

This work was supported by the National Key Research and Development Program (2024YFA1803400), the Strategic Priority Research Program of the Chinese Academy of Sciences (Grant Nos. XDB1480000, XDB0520301), the National Natural Science Foundation of China (Grant Nos. 22177128, 22407137), the Natural Science Foundation of Jiangsu Province (Grant Nos. BK20243004, BK20232046), the Science and Technology Project of Suzhou (Grant No. SYG2024080). We thank Prof. Mao Lanqun (College of Chemistry, Beijing Normal University) for FSCV study, Prof. Xu Guangyin (Institute of Neuroscience, Soochow University) for PD model’s EMG monitoring, Suzhou NIR-Optics Technology Co., Ltd. for its instrumental and technique support on the fluorescence imaging.

## AUTHOR CONTRIBUTIONS

Conceptualization: Wang Q.B., Chen G.C., Yin H.Q.; Methodology: Yin H.Q., Li T.W., Jiang W.Q.; Investigation: Yin H.Q., Li T.W., Jiang W.Q., Wu F., Yuan Q.H., Wang T.; Visualization: Yin H.Q.; Funding acquisition: Wang Q.B., Chen G.C.; Project administration: Wang Q.B., Chen G.C., Zhang Y.J.; Supervision: Wang Q.B., Chen G.C., Li C.Y., Jiang J.; Writing-original draft: Yin H.Q., Chen G.C.; Writing-review & editing: Wang Q.B., Chen G.C., Jiang J.

## DECLARATION OF GENERATIVE AI AND AI-ASSISTED TECHNOLOGIES IN THE WRITING PROCESS

During preparation of this paper, the authors did not use AI tools or services.

## STAR METHODS

### KEY RESOURCES TABLE

| REAGENT OR RESOURCE | SOURCE | IDENTIFIER |
| --- | --- | --- |
| **Antibodies** |  |  |
| Mouse monoclonal TRPV1 antibody | Abcam | Cat#ab203103 |
| Mouse monoclonal HaloTag antibody | Promega | Cat#G9211 |
| Rabbit Monoclonal Flag antibody | CST | Cat#14793 |
| Mouse Monoclonal Flag antibody | CST | Cat#8146 |
| Rabbit Monoclonal c-Fos antibody | CST | Cat#31254 |
| Rabbit Monoclonal IBA-1 antibody | CST | Cat#17198 |
| Rabbit Monoclonal Tyrosine Hydroxylase antibody | CST | Cat#58844 |
| Anti-Rabbit Alexa Fluor<sup>®</sup> 488 Conjugate | CST | Cat#4412 |
| Anti-Mouse Alexa Fluor<sup>®</sup> 647 Conjugate | CST | Cat#4414 |
| **Chemicals** |  |  |
| DSPE-PEG<sub>2000</sub>-NHS | Ruixibio | Cat#R-0042 |
| MPTP hydrochloride | MCE | Cat#HY-15608 |
| Calbryte 630AM | AAT Bioquest | Cat#20721 |
| Hoechst 33342 | Beyotime | Cat#C1022 |
| Lipo6000 Transfection Reagent | Beyotime | Cat#C0526 |
| OCT compound | Tissue-Tek | Cat#4583 |
| **Experimental models: Cell lines** |  |  |
| HEK 293T cells | NCACC | Cat#GNHu44 |
| **Experimental models: Organism/strains** |  |  |
| C57BL/6J | GemPharmatech | Cat#000013 |
| **Recombinant DNA** |  |  |
| pCDNA3.1-CMV-HaloTRP-3×Flag-t2a-EGFP | This paper | N/A |
| pCDNA3.1-CMV-TRPv1-3×Flag-t2a-EGFP | This paper | N/A |
| **Bacterial and virus strains** |  |  |
| AAV-hSyn-HaloTRP-3×Flag | This paper | N/A |
| AAV-hSyn-TRPV1-3×Flag | This paper | N/A |
| **Software and algorithms** |  |  |
| AlphaFold | DeepMind | https://alphafoldserver.com/ |
| Pymol 3.1 | Schrödinger, Inc. | https://www.pymol.org/ |
| AutoDockFR | Sanner et al. | https://ccsb.scripps.edu/adfr/ |
| ImageJ | Rasband et al. | https://imagej.nih.gov/ij/ |
| Neuroexplorer | Nex Tech. | https://www.neuroexplorer.com/ |
| Offline Sorter | Plexon | https://plexon.com/products/offline-sorter/ |
| Originlab 2024b | OriginLab Corp. | https://www.originlab.com |
| IBM SPSS statistics 22 | IBM | https://www.ibm.com/software |

### METHOD DETAILS

#### Synthesis of HPN

Firstly, a NIR-II photothermal conversion molecular (NPCM) with a D-A-D structure was synthesized according to the previous study with some changes (Figure S2A). ^1^H NMR (400 MHz, DMSO-d6): δ 7.95-7.84 (m, 1H), 7.82-7.68 (m, 4H), 7.65-7.56 (m, 4H), 7.53 (d, J = 17.0 Hz, 8H), 7.11 (d, J = 8.5 Hz, 4H), 6.53-6.26 (m, 2H), 4.89 (d, J=2.4 Hz, 4H), 3.52 (s, 2H), 3.15-2.96 (m, 4H), 2.82-2.73 (m, 4H). ESI-MS (M-BF_4_) m/z: calcd for C_49_H_37_O_2_S^2+^, 721.2229; found, m/z 721.2229. Next, a NIR-II photothermal conversion nanomicelle (PN) was prepared. NPCM and DSPE-PEG_2000_-NHS were co-dissolved at a mass ratio of 1:20 in *N*,*N*-dimethylformamide (DMF), which was added dropwise into deionized water (1:100, volume ratio) under stirring with 1000 rpm to self-assemble into a homogeneous PN solution. Subsequently, the micelles solution was adjusted to mild alkalinity to pH 8.3-8.5 for the reaction of NHS and the HaloTag chloroalkane ligands (HTL). HTL (molar ratio of DSPE-PEG_2000_-NHS: HT = 1:1) was dissolved in DMF, which was added dropwise into the PN solution under stirring (500 rpm) overnight. Finally, PN was functionalized with chloroalkane to obtain HPN with self-labeling capacity.

#### Characterizations of HPN

The morphology of HPN was characterized using a Tecnai G2 F20 S-Twin transmission electron microscopy (TEM, FEI, USA) operated at an acceleration voltage of 200 kV. The hydrodynamic diameter and surface charge were measured by a Zetasizer Nano ZS Malvern Nanosizer. The UV-Vis-NIR absorption spectrum of HPN was measured using a Perkin-Elmer Lambda 25 UV-Vis spectrometer. The NIR emission spectrum was recorded using an Applied NanoFluorescence spectrometer (USA) with 785 nm excitation. The photoacoustic properties of HPN were examined by a LOIS-3D Optoacoustic Imaging System (TomoWave Systems, Houston, TX, USA) equipped with a wavelength tunable laser (660 nm-2300 nm) and a pulse duration of 8 ns.

#### Photothermal conversion effect and conversion stability of HPN

HPN dispersed in PBS (50 μg/mL, 1 mL) was placed into a quartz cuvette (3.5 mL) and illuminated using a 1064 nm laser (Changchun New Industries Optoelectronics Tech. Co., Ltd.) at a power density of 1.0 W/cm^2^. The temperature during the photothermal process was monitored in real time by a dual-input J/K digital thermometer (DM6801) during both heating and cooling phases. To assess the photothermal conversion stability of HPN, repetitive irradiation was used to measure the photothermal effect at 1 day, 1 month and 2 months after HPN preparation.

#### Calculation of photothermal conversion efficiency

Photothermal conversion efficiency (PCE) of HPN aqueous solution (50 μg/mL) was calculated according to the previous study^68^, and water was used as control. Equation of PCE was as follows:

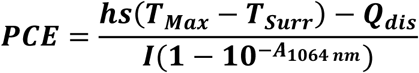

Where *h* is the heat transfer coefficient, *S* is the surface area of the container, *T_Max_* and *T_Surr_* respectively represented the maximum temperature of photothermal process and ambient temperature (27 ℃). *A_1064 nm_* is the absorption intensity of solution at 1064 nm. The value of *hs* was evaluated from following Equation:

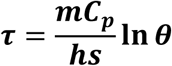

Where *θ* is calculated as (*T* – *T_Surr_*)⁄(*T_Max_* − *T_Sure_*), *τ* is the time of cooling phase, *m* is the mass of the solution (1 g). *C_p_* represents the specific heat capacity of the solvent (4.2 J g^-1^ ℃^-1^).

*Q_dis_* is heat dissipation from laser mediated by solvent and container, which was calculated by Equation as follows:

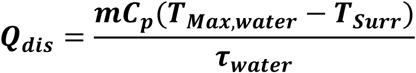

Where *τ_water_* and *T_Max, water_* were determined to be 306 and 28.8 ℃.

#### Molecular docking and visualization

The core segments of the monomer HaloTRP include the TRPV1-S1, S1-S2 loop, cpHaloTag, and TRPV1-S2 subunits, and their amino acid sequences were as follows: IFYFNFFVYCLYMIIFTTAAYYRPVEGLPPYKLNNTVGDFARETFQAFRTTDVGRKLIIDQNVFIEGTLPMGVVRPLTEVEMDHYREPFLNPVDREPLWRFPNELPIAGEPANIVALVEEYMDWLHQSPVPKLLFWGTPGVLIPPAEAARLAKSLPNCKAVDIGPGLNLLQEDNPDLIGSEIARWLSTLEISGEPTTGGSGGTGGSGGTGGSMAEIGTGFPFDPHYVEVLGERMHYVDVGPRDGTPVLFLHGNPTSSYVWRNIIPHVAPTHRCIAPDLIGMGKSDKPDLGYFFDDHVRFMDAFIEALGLEEVVLVIHDWGSALGFHWAKRNPERVKGIAFMEFIRPIPTWDEWPEFAGYFRVTGEILSVSGGVYFFFRGIQYFL

The 3D structures of HaloTRP_m_ and the tetramer HaloTRP (HaloTRP) were modeled using Alphafold Server (<u>alphafoldserver.com</u>), and visualization of 3D models was performed using Pymol 3.10. ChemBioDraw Ultra 14.0 was used for drawing the 2D structure of chemical molecules, including the HaloTag ligand, while ChemBio3D Ultra 14.0 was used for converting the 2D structures to 3D models. Docking of HTL to the cpHaloTag domain of HaloTRP was performed using AutoDockFR.

#### Plasmids and adeno-associated virus (AAV) of HaloTRP construction

The plasmids we used in our study included pCDNA3.1-CMV-HaloTRP-3×Flag-t2a-EGFP and pCDNA3.1-CMV-TRP-3×Flag-t2a-EGFP. AAV was used for neuron transduction *in vivo*, and the AAV used in this study included AAV5-hSyn-HaloTRP-3×Flag and AAV5-hSyn-TRP-3×Flag. It’s suggested that the promoter of synapsin (Syn) specifically introduces target gene expression into neurons without expression in glia. The plasmids and AAVs were all constructed and packaged by GeneChem Co. Ltd. The plasmid of HaloTRP can be accessed with Addgene (ID: 231245).

#### Cell culture and transfection

HEK 293T cells were purchased from the National Collection of Authenticated Cell Cultures of China. The cells were cultured in Dulbecco’s modified Eagle’s medium (DMEM, Gibco, Invitrogen) supplemented with 10% fetal bovine serum (FBS, Gibco, Invitrogen) at 37 ℃ in a 5% CO_2_/95% air incubator. Lipo6000 Transfection Reagent (Beyotime, Shanghai, China) was applied for transfection of HaloTRP and TRP plasmids according to the manufacturer’s instructions.

#### HPN specifically labels HaloTRP cells *in vitro*

After HaloTRP or TRP transfection with HEK 293T cells, HPN (50 μg/mL) diluted in DMEM was incubated with the HaloTRP and TRP cells, respectively, for 4 h. Then the cells were washed with PBS for three times, followed by imaging with an Olympus IX71 fluorescence microscope (Olympus Corporation, Tokyo, Japan). The transfected cells were imaged at 488 nm with GFP fluorescence, and the fluorescent signal of HPN was detected at 555 nm.

#### Calcium imaging

Calcium imaging was used to detect NIR-II sensitivity of HaloNeu. After cells were treated with HaloNeu, the calcium ion indicator Calbryte 630 AM (10 μM, AAT Bioquest) diluted in DMEM was incubated with cells for 30 min at 37 ℃. After incubation, the cells were washed with PBS three times. Then the Olympus IX71 fluorescence microscope was used for calcium imaging under 1064 nm irradiation (1 W/cm^2^, 3 s), and the images were recorded every 0.5 s for 10 s with excitation of 608 nm and emission of 622 nm. The fluorescent signal of Calbryte 630 AM could be completely distinguished from GFP and HPN. Image intensity was calculated using ImageJ. Moreover, real-time calcium imaging was also performed with 1064 nm at 60 mW/cm^2^ for 5 min stimulation, with images recorded every 1 s. Calcium fluctuations of unlabeled TRP cells (TRP-HPN) were also tested under 1064 nm irradiation.

#### Whole-cell patch-clamp recording

We employed the whole-cell patch-clamp technique to investigate the modulatory effect of 1064 nm light-mediated activation of HaloNeu on cellular electrical activity. The patch-clamp system (HEKA, Munich, Germany) was equipped with an upright multi-channel fluorescence microscope (Suzhou NIR-Optics Technology Co., Ltd., China) for locating the HaloTRP cells (GFP tagged, 488 fluorescence) labeled with HPN (555 nm). Moreover, a 1064 nm laser (Changchun New Industries Optoelectronics Tech. Co., Ltd.) was also configured on the excitation light path for NIR-II stimulation, and the switching of the 1064 nm laser was triggered by a TTL pulse that was programmed and outputted by an EPC 10 Double USB Amplifier (HEKA, Munich, Germany). Firstly, HaloNeu cells were searched by multi-channel fluorescence microscope, and patched using the whole-cell current-clamp mode. After the cells were stabilized for 5 min, the membrane potential was recorded by the EPC10 Double USB Amplifier with a resting holding potential of -60 mV. Switching of 1064 nm laser (1 W/cm^2^, 3 s, 5 circles) were triggered. Membrane potential of HaloNeu cells were recorded only when the seal resistance was > 500 MΩ, and the series resistance (< 30 MΩ) changed < 20% throughout the experiment. For membrane potential recording, the bath solution was contained (in mM): 140 NaCl, 5 KCl, 2 CaCl_2_, 1 MgCl_2_, 10 HEPES, and 10 D-glucose, adjusted to pH 7.3 with NaOH, 300∼310 mOsm. The internal solution was contained (in mM): 125 KCl, 5 NaCl, 2.5 CaCl_2_, 2 MgCl_2_, 5 EGTA, 10 D-glucose, and 5 HEPES adjusted to pH 7.3 with KOH, 295∼300 mOsm. Additionally, membrane potential changes in HaloNeu cells with ultralow power of 1064 nm (60 mW/cm^2^, 5 min) were also recorded by whole-cell patch-clamp technique. The control experiment using TRP-HPN cells was also performed under same conditions.

#### Stereotaxic injection of AAV virus and HPN *in vivo*

Adult (6 weeks old) male C57BL/6J mice were used for the animal subjects in this study. All animal operations were performed according to the Chinese Regulations for the Administration of Affairs Concerning Experimental Animals, China (GB/T 35892-2018) and approved by the Animal Ethics Committee of Suzhou Institute of Nano-Tech and Nano-Bionics, Chinese Academy of Sciences, with the animal ethics approval number SINANO-2023-020. Mice were group-housed at 23 ℃ on a cycle of 12-h light: 12-h dark cycle at a relative humidity of 40-48% and free access to food and water. All efforts were made to reduce the number of mice sacrificed and to alleviate their suffering. Mice were anesthetized with intraperitoneal injection of sodium pentobarbital (50 mg/kg) and placed on the stereotaxic frame (RWD Life Science, Shenzhen, China). VTA region was chosen for AAV virus injection, and the stereotaxic coordinate was as follows: ±0.5 mm lateral to the middle (ML: ±0.5 mm), 3.5 mm posterior to bregma (AP: -3.5 mm) and 4.5 mm beneath the dura (DV: -4.5 mm). AAV5-hSyn-HaloTRP-3×Flag (1.27*10^12^ v.g./mL, 1.0 μL) was injected into both sides of VTA region using a Hamilton 10 µL syringe equipped with a 33 g needle, and the injection process of syringe was driven by a syringe pump (KDS Legato 100, KD Scientific, USA) at the speed of 0.1 μL/min. After injection, the needle was kept in the brain for 10 min to avoid reflux. The wound was sealed with degradable sutures and applied to antibiotic ointment. After recovery, mice were returned to cage and free access to food and water. Three weeks after AAV virus injection, HPN (1.0 mg/mL, 1.0 μL) was stereotaxic injected at the same site. The procedures of injection were similar with that of AAV virus.

#### Histological analysis

After treatment, the anesthetized C57BL/6J mice were perfused with PBS, and brain tissues were sampled and embedded in OCT compound (Tissue-Tek, Sakura Finetek Japan Co., Japan) without fixation. Sections of frozen brain slices with a thickness of 15 μm were obtained using a pathological slicer at -20 ℃ (Leica CM 3050S, Leica Biosystems). The slices were washed with PBS three times for 10 min, and permeabilized with 0.1% Triton X-100 (Beyotime, Shanghai, China) for 15 min, followed by 5% normal goat serum (Solarbio, Beijing, China) blocking for 1 h. Then the blocked slices were incubated with primary antibodies solution overnight. After incubation, slices were washed with PBS three times for 10 min, and secondary antibodies were applied to the slices for 1 h. Finally, the nuclei were stained with Hoechst 33342 (Beyotime Life Science, Shenzhen, China) for 5 min, after which the fluorescence of the whole-brain slices were scanned by Leica DMi8 Widefield microscope with Leica Application Suite X (LAS X) software. The primary antibodies were as follows: anti-TRPV1 (1:1000, ab203103, Abcam), anti-HaloTag (1:500, G9211, Promega), anti-Flag (1:1000, #14793, CST), c-Fos (1:1000, #31254, CST), IBA-1 (1:500, #17198, CST), Tyrosine Hydroxylase (1:1000, #58844, CST). Secondary antibodies included Anti-rabbit Alexa Fluor^®^ 488 Conjugate (1:1000, #4412, CST), Anti-rabbit Alexa Fluor^®^ 647 Conjugate (1: 1000, #4414, CST).

#### Photoacoustic (PA) imaging

The Laser Optoacoustic Imaging System LOIS-3D (TomoWave Systems, Houston, USA) equipped with a broad wavelength laser (660-2300 nm) was used for PA characterization of HPN at different concentration in PBS. For PA signal detection of HPN in the brain, the Laser Optoacoustic Imaging System Maestro HR-M (TomoWave Systems, Houston, USA) was equipped with an unfocused linear probe. The probe was placed onto the brain of the mouse for scanning the PA signal, and the gap between the probe and the brain was filled with ultrasonic coupling agent.

#### Establishment of chronic Parkinson’s disease (PD) model

Metabolism of MPTP in the brain to the neurotoxin MPP^+^ specifically damages dopaminergic neurons in the pars compacta of substantia nigra, causing permanent Parkinson’s disease symptoms in humans and non-human primates. The adult male C57BL/6J mice were intraperitoneally injected with MPTP-HCl (30 mg/kg, M0896, Sigma) every two days for three consecutive weeks. Mouse body weight was recorded every other day. Finally, histological analysis was used to detect the significant decrease in DA neurons in the STR and SNc etc. for PD model determination.

#### *In vivo* electrophysiological recording

After HaloNeu was delivered into VTA region, the mice were anesthetized and secured in stereotaxic frame, placed on a heating blanket. Microwire array electrodes with 16 channels (Kedou BC, Suzhou, China) were acutely implanted into VTA region (AP: - 3.5 mm; ML: ±0.5 mm; DV: -4.5 mm), and the *in vivo* electrophysiological activity was recorded by 256-channel Neurolego recording system (Greatink, Nanjing, China) with a 0.3-10 kHz sorting unit for the firing spikes acquired at 20 kHz sampling rate. After the baseline was recorded for 1 min, continuous 1064 nm light (1 W/cm^2^, 60 mW/cm^2^) illuminated mouse brain from a distant collimator connected to a 1064 laser for 5 min. The electrophysiological activity was recorded for 10 min. Recording was performed at different time points after HaloNeu delivery 1 day, 1 week, 2-week, 1 month and 2-month. Data analysis was performed using NeuroExplorer and Offline Sorter.

#### Behavioral activity test with open field test

After HaloNeu was stereotaxically delivered into the VTA region of mice, the open field test was used to measure behavioral activity of mice with 1064 nm irradiation. The mice were placed individually in a corner of the open field composed of white acrylic panels (30 × 30 × 30 cm^3^). The open field was divided into 16 grids, and the center 4 grids belonged to the central area. The behavioral activity of test mice was monitored by a camera and analyzed using SuperMaze software (Xinruan, Shanghai, China). Firstly, the mice were habituated to the testing open field for 30 minutes prior to the experiment. Each mouse was tested for 5 min without 1064 nm irradiation, and the metabolites and odor were eliminated with 75% ethanol after each test was finished. For the pairing study in the open filed, the interval time between two tests was more that 1 h to ensure the recovery of the mice. After a break, the behavioral activity of previously tested mice was performed in the open field under 1064 nm irradiation for 5 min. A beam expander was installed 35 cm above the center of the open field and connected to a 1064 nm laser at a working power of 20 W. The mean power density of the 1064 nm laser illuminating the mice was calculated to be about 60 mW/cm^2^. The open field test was performed at different time points (1 day, 1 week, 2-week, 1 month and 2 months) after HaloNeu delivery. For the behavioral activity measurement of PD mice treated with the HaloNeu approach, the procedures were followed those described above.

#### Fast-scan cyclic voltammetry (FSCV) *in vivo*

After HaloNeu was delivered into VTA region, FSCV was used to real-time detect the concentration of DA neurotransmitter released from synapses of VTA DA neurons at different time points (1 day and 2 months) after HPN injection. FSCV *in vivo* was performed as previously reported. In brief, a carbon fiber microelectrode (CFME) was implanted into the NAc region where synapses of VTA DA neurons project. Ag/AgCl wire was used as a reference electrode and positioned on the dura of the brain. An Ag wire fixed to the brain skin acted as a grounding electrode. A Patch-clamp Amplifier (HEKA, EPC10, Germany) was used to run a cyclic voltammogram (CV) with a triangular waveform potential (-0.5 mV to +1.1 mV vs. Ag/AgCl) at a scan rate of 400 V/s, repeated every 300 ms, and the sampling rate was 200 kHz. Patchmaster software was used to collect electrochemical data. All electrodes were cycled for at least 15 min to stabilize the background currents. After stabilization, background currents were subtracted from at least ten successive CVs before each recording. A 1064 nm irradiated (60 mW/cm^2^) the mouse brain for 90 s after a 1 min baseline recording. Every CFME was calibrated with a standard DA solution *in vitro* before recording, and a curve of concentration vs. current was plotted, which was used for converting between current and concentration.

#### Electromyography (EMG) analysis of PD mouse

EMG of muscle activity is considered the gold standard for the diagnosis of tremor. The gastrocnemius in the hindleg of PD mice suffers from dysfunction including pain and cramping, which is closely related to tremor symptoms. For PD tremor monitoring, the gastrocnemius was the focus of EMG recording in our study. A tinned copper terminal wire (30 AWG, Skygod) was fixed on the gastrocnemius, acting as a recording electrode of EMG. Another wire without contacting the recording electrode was fixed on an adjacent muscle. The wires passed under the back skin and were fixed to a head-plate without affecting mouse behavior. EMG signal was collected using a Biopac System MEC 100 Amplifier with a filter of 300 Hz (Biopac System Inc., Holliston, MA). After the EMG baseline of the PD mouse was recorded for 5 min, a 1064 nm laser from a distant collimator connected to a 1064 nm laser illuminated the PD mouse (60 mW/cm^2^, 5 min). The duration of EMG recording was 15 min. The area of EMG curve was analyzed by AcqKnowledge software. EMG signals from 5 min baseline (1064 nm pre-irradiation), 5 min of 1064 nm irradiation and 5 min post-irradiation were calculated. EMG signals of the PD HaloNeu mice were recoded at time points of 1 day, 1 week, 2 weeks, 1 month, and 2 months after HaloNeu delivery.

#### Statistical analysis

All quantitative data are presented as the Mean ± S.D. Comparisons of the fluorescent colocalization coefficient, PA intensity, HPN characteristics were performed using one-way ANOVA for significance analysis of multiple samples, followed by post-hoc correction with the LSD test. The data including the open field test, electrophysiological recording, and EMG were analyzed using a two-tailed paired sample t-test. Other statistical analyses were performed using an independent sample t-test. All statistical analyseses were performed with the IBM SPSS statistics 22 software, and a P-value of < 0.05 was considered significant. All data were from at least 3 replicates.

