## Supplemental Figures S1-S7 and Supplemental Table S1 for "Near-Infrared-II Chemo-Optogenetics for Deep-Brain Stimulation"

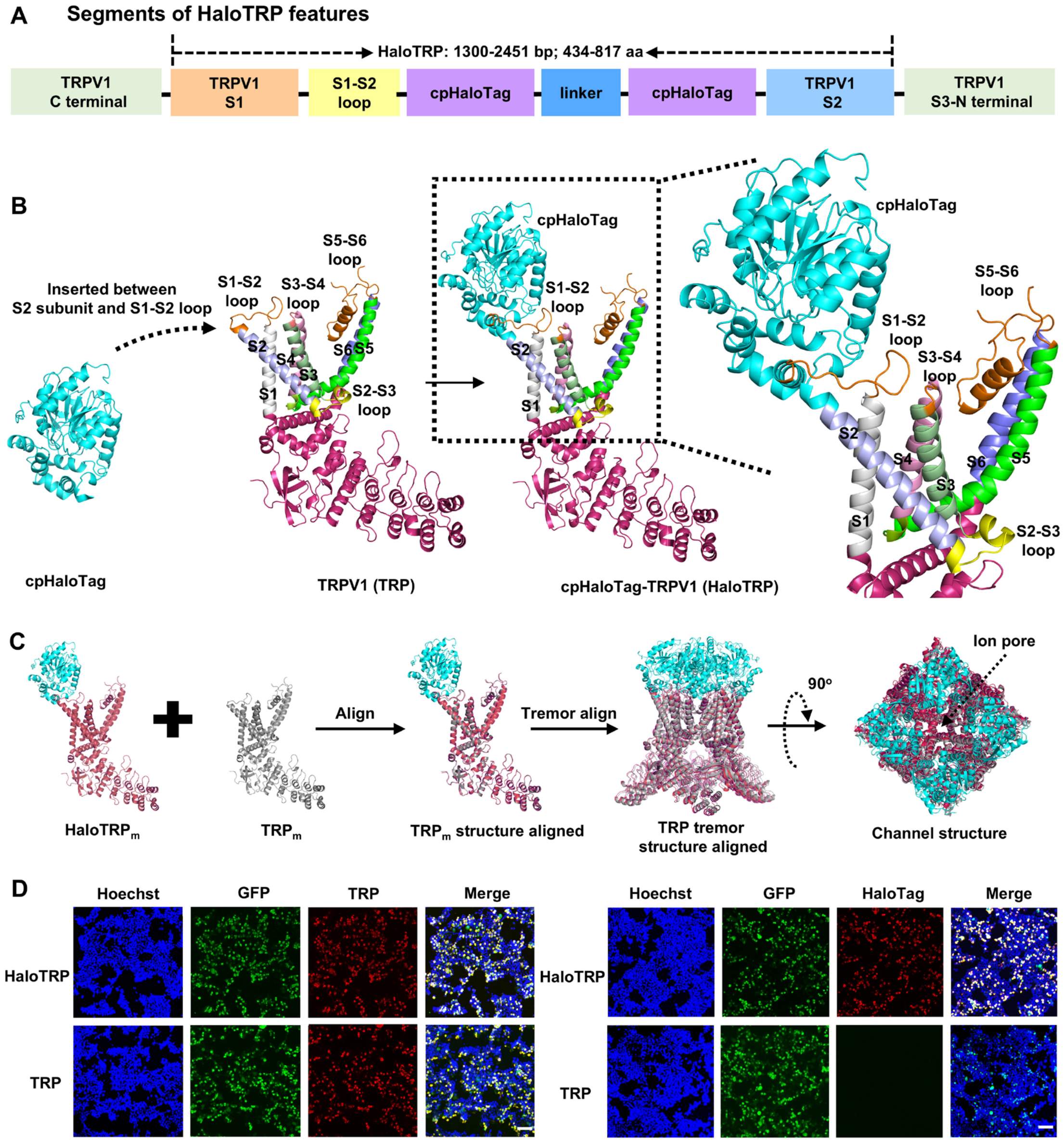


**Figure S1. Design and expression of HaloTRP. (A)** Design and core segments of HaloTRP. **(B)** The 3D structure of $HaloTRP_m$ formation. cpHaloTag was inserted between S2 subunit and S1-S2 loop of TRPV1. **(C)** Structural alignment analysis of HaloTRP and TRP, suggesting the preservation of TRP structure in HaloTRP. **(D)** Immunofluorescence imaging of TRP and HaloTag in 293T cells expressing HaloTRP, confirming co-expression of both TRP and HaloTag. Scale bar, 100 μm.

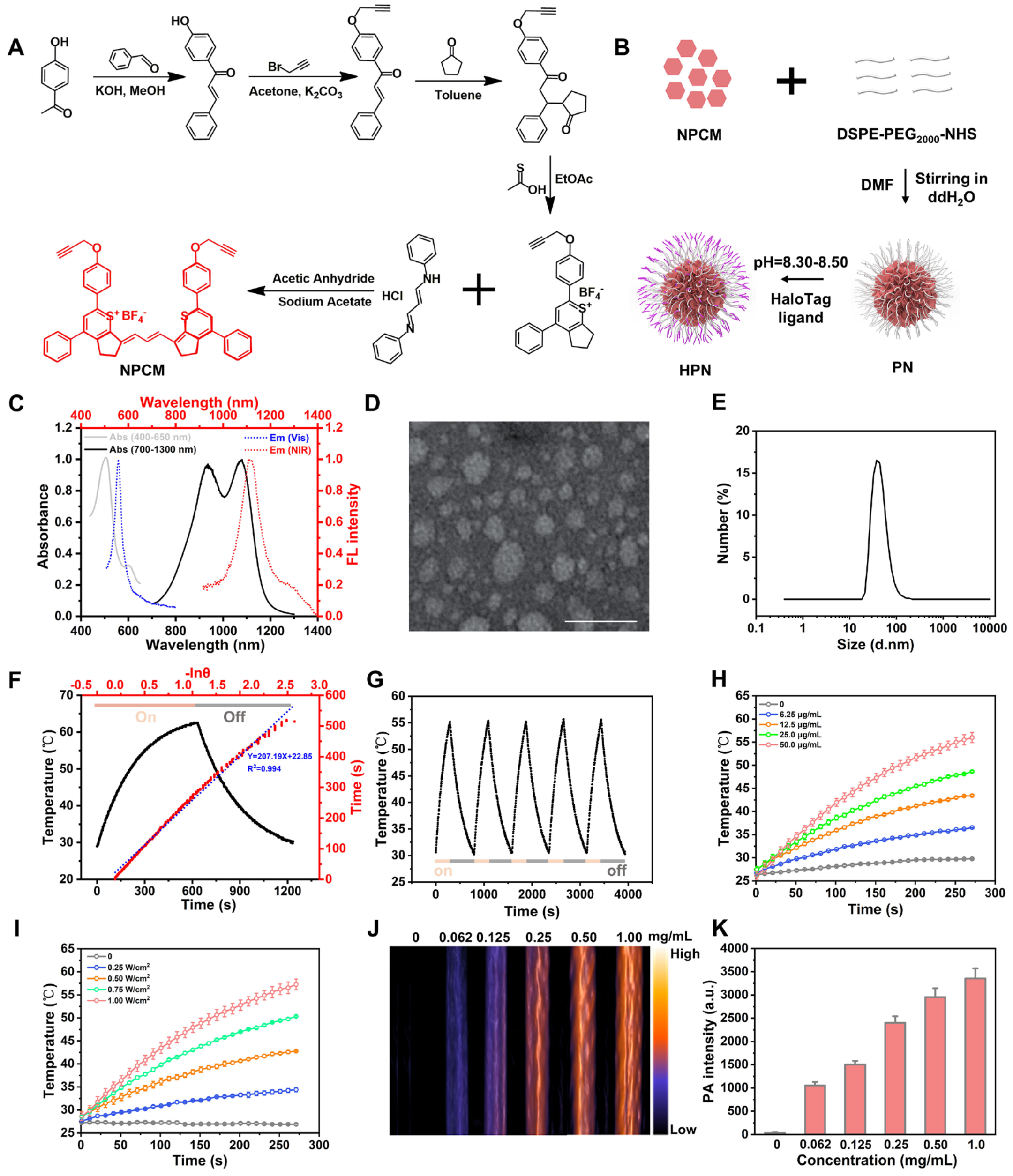


**Figure S2. Preparation and characterizations of HPN. (A)** Process of NPCM synthesis. **(B)** Schematic of HPN preparation. **(C)** Absorption and emission spectra of HPN. **(D)** TEM image of HPN. Scale bar, 50 nm. **(E)** DLS profile of HPN. **(F)** Photothermal conversion efficiency of HPN (50 μg/mL) under 1064 nm laser irradiation (1 W/cm$^2$) showing a high photothermal conversion efficiency of 71%. **(G)** Photothermal stability detection of HPN (50 μg/mL) over multiple on/off cycles of 1064 nm laser irradiation (1 W/cm$^2$). **(H)** Photothermal response of HPN at varying concentrations under 1064 nm irradiation (1 W/cm$^2$, n=3). **(I)** Temperature elevation of HPN (50 μg/mL) as a function of 1064 nm laser power density (n=3). **(J and K)** Photograph of HPN PA signal at different concentrations. Statistical data were presented as Mean ± S.D.

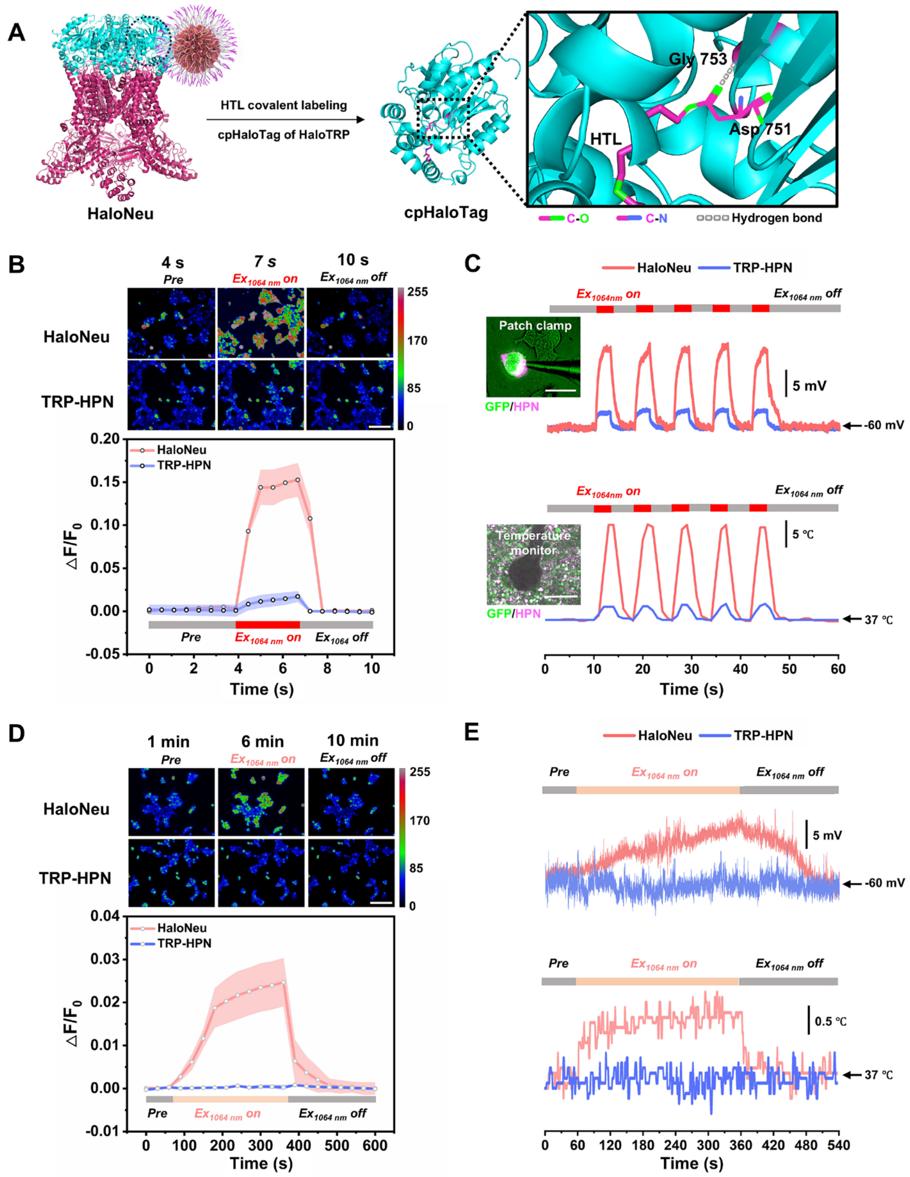


**Figure S3. 1064 nm illumination induces calcium ion influx and depolarize membrane potential in HaloNeu cells. (A)** Asp751 of HaloTRP forms an irreversible ester bond with the chloroalkane ligand HTL of HPN, while Gly753 participates in stabilizing the complex through hydrogen bonding. **(B)** Calcium imaging reveals an increase in intracellular calcium ion concentration in HaloNeu-expressing cells under continuous 1064 nm irradiation (1 W/cm$^2$, 3 s). Scale bar, 100 μm. **(C)** HaloNeu mediates sustained membrane potential depolarization in response to 1064 nm light with rapid temperature changes (1 W/cm$^2$, 3 s, 5 cycles). Scale bar (Patch clamp), 20 μm. Scale bar (Temperature monitor), 50 μm. **(D)** A gradual increase in HaloNeu cells and minimal response in TRP-HPN cells show that ultralow power of 1064 nm irradiation (60 $mW/cm^2$) sufficiently activates HaloNeu. Scale bar, 100 μm. **(E)** Representative traces of membrane potential in HaloNeu and TRP-HPN cells under low power of 1064 nm irradiation with a slight temperature increase are show. Statistical data were presented as Mean ± S.D.

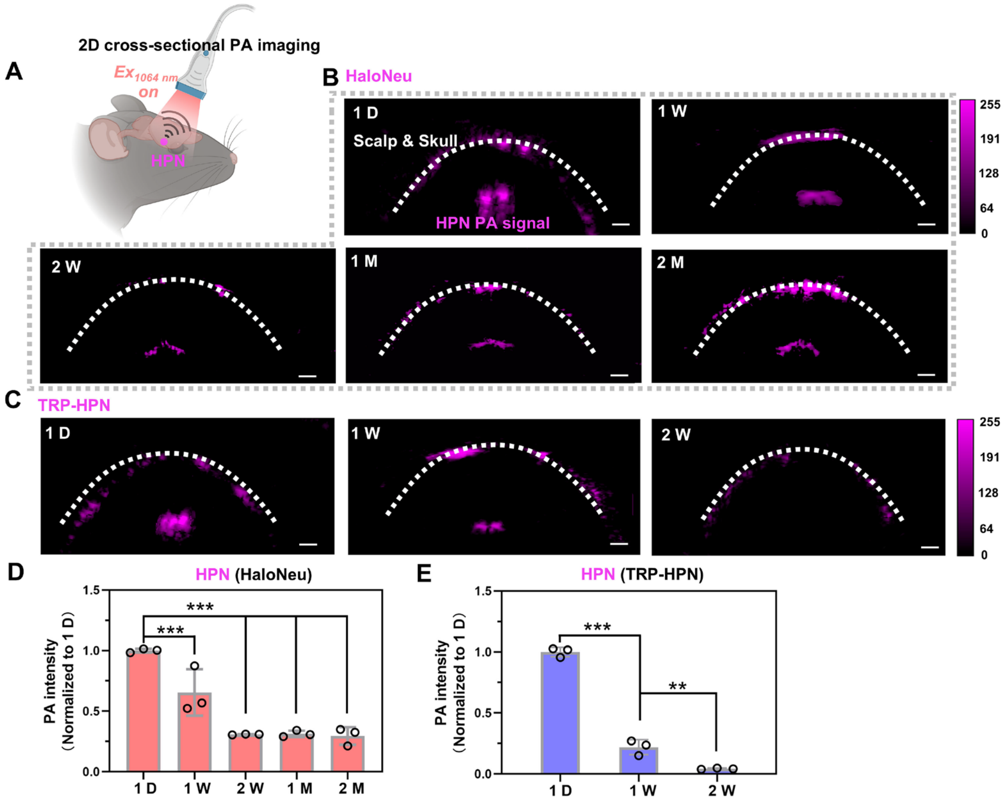


**Figure S4.** ***In vivo*** **PA imaging of HPN. (A)** Schematic diagram of 2D cross-sectional PA imaging of the mouse brain. **(B)** PA imaging of HPN in the HaloNeu mouse brain, showing persistent PA signal from 1 day to 2 months post-injection. PA signals were consistently detected at a depth of approximately 4.5 mm beneath the skull within the VTA region. Scale bar, 1 mm. **(C)** 2D PA imaging of HPN signals in TRP-HPN mouse brain. Scale bar, 1 mm. **(D and E)** Statistical analysis of PA signal intensity in HaloNeu **(D)** and TRP-HPN **(E)** mice over time (n=3). HPN PA signals exhibited their long-term retention in HaloNeu mouse for up to 2 months. Statistical data were presented as Mean ± S.D.

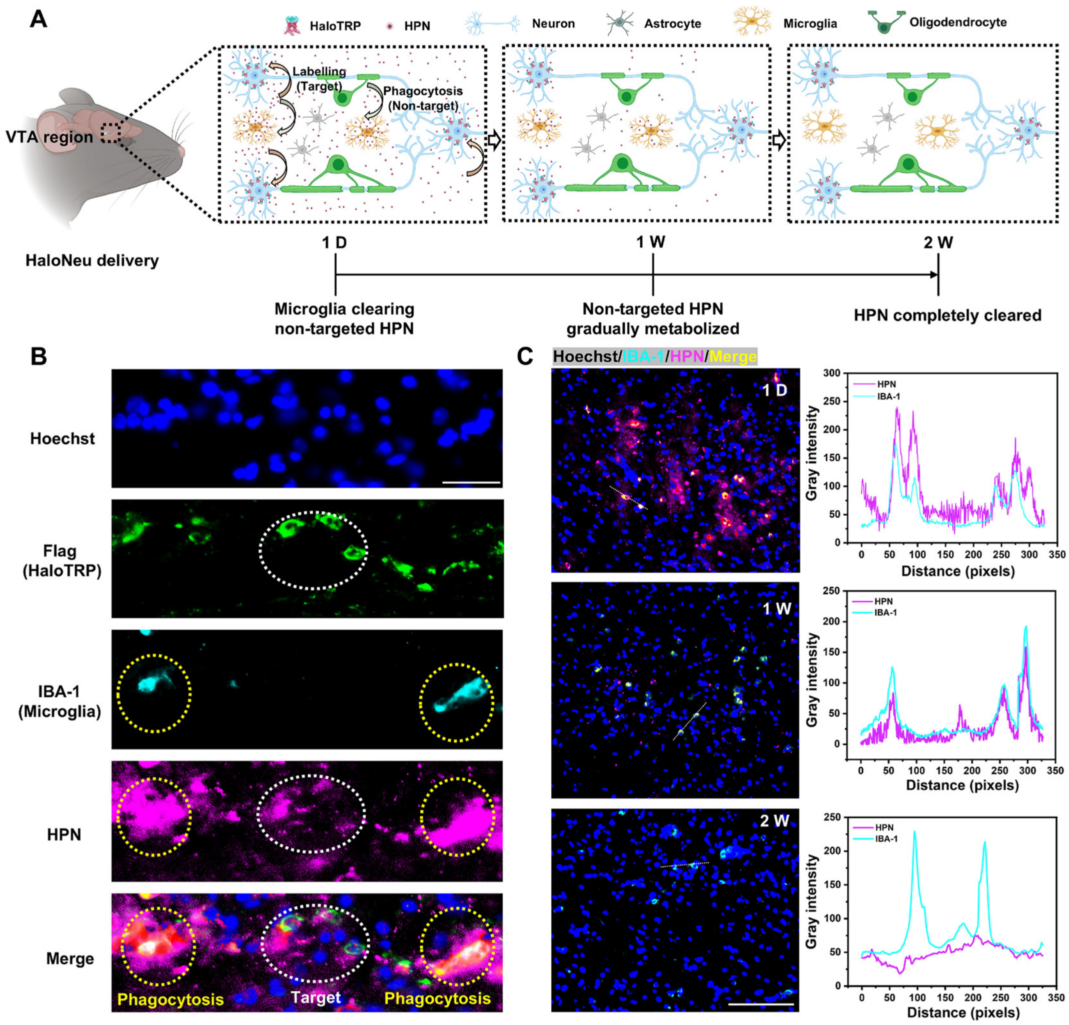


**Figure S5. Microglia clearance of untargeted HPN within 2 weeks. (A)** Schematic diagram of microglial clearance of material non-targeted HPN, with complete clearance within 2 weeks. **(B)** Immunofluorescence imaging showing IBA-1-positive microglia participate in HPN clearance in HaloNeu mice. Scale bar, 25 μm. **(C)** Complete clearance of untargeted HPN from brain tissue within two weeks in HaloNeu mice. Scale bar, 100 μm.

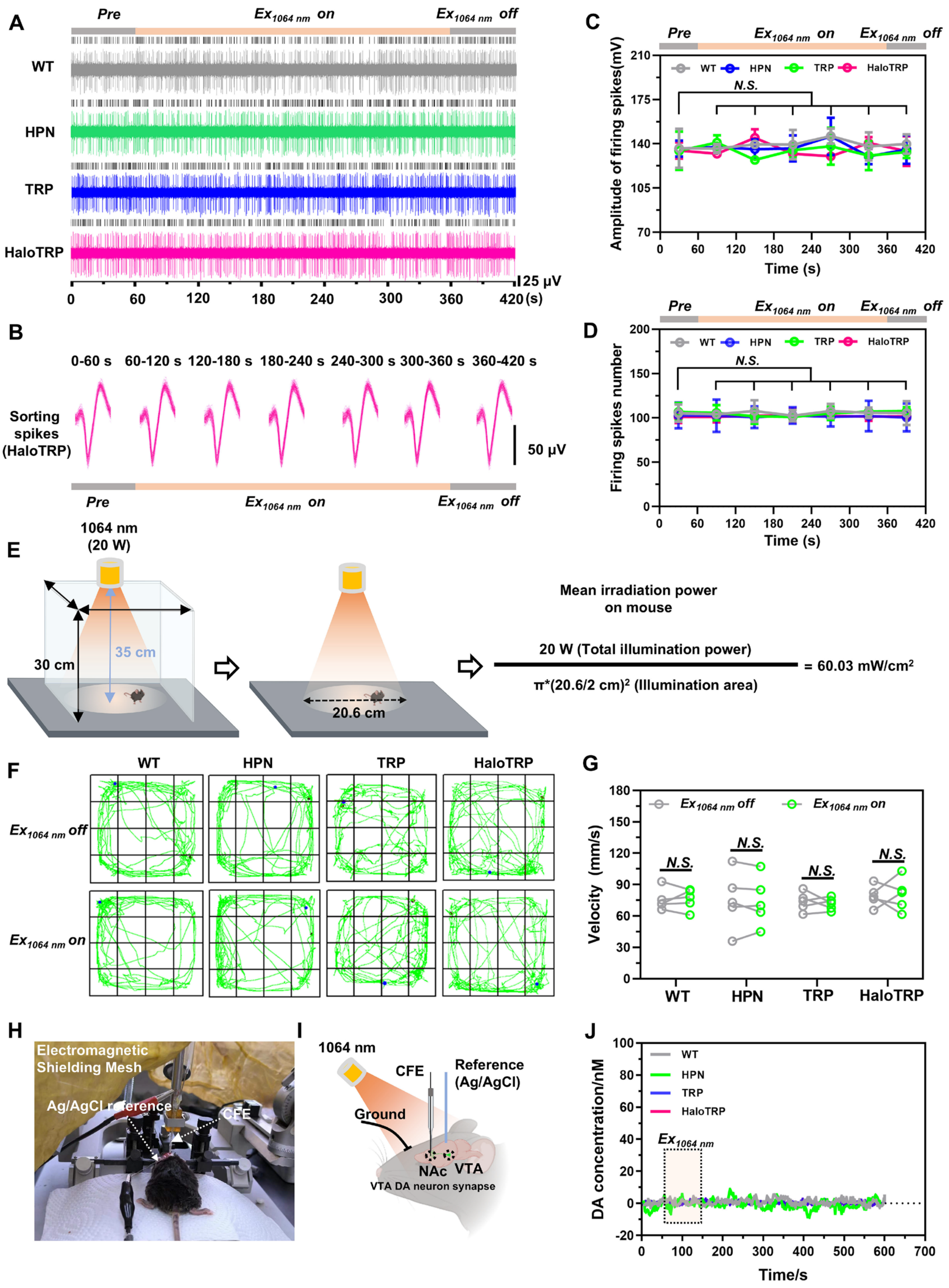


**Figure S6. Firing spikes, behavioral activity and DA concentration in mice treated with HPN, TRP or HaloTRP alone showed no significant changes under 1064 nm illumination. (A)** Neural firing spikes in VTA region under 1064 nm irradiation across groups. **(B)** Sorted firing spikes of VTA neurons in HaloTRP mouse across different time periods, before, during, and after 1064 nm illumination. **(C and D)** Statistical analysis of firing spike counts **(C)** and amplitude **(D)** across groups (n=3). **(E)** Determination of mean power density for 1064 nm irradiation in mice. **(F)** Behavioral activity across groups in the absence or presence of 1064 nm illumination. **(G)** Quantification of locomotor velocity crossing the center grid across groups (n=5). Statistical data were presented as Mean ± S.D. $P \geq 0.05$ *N.S.*

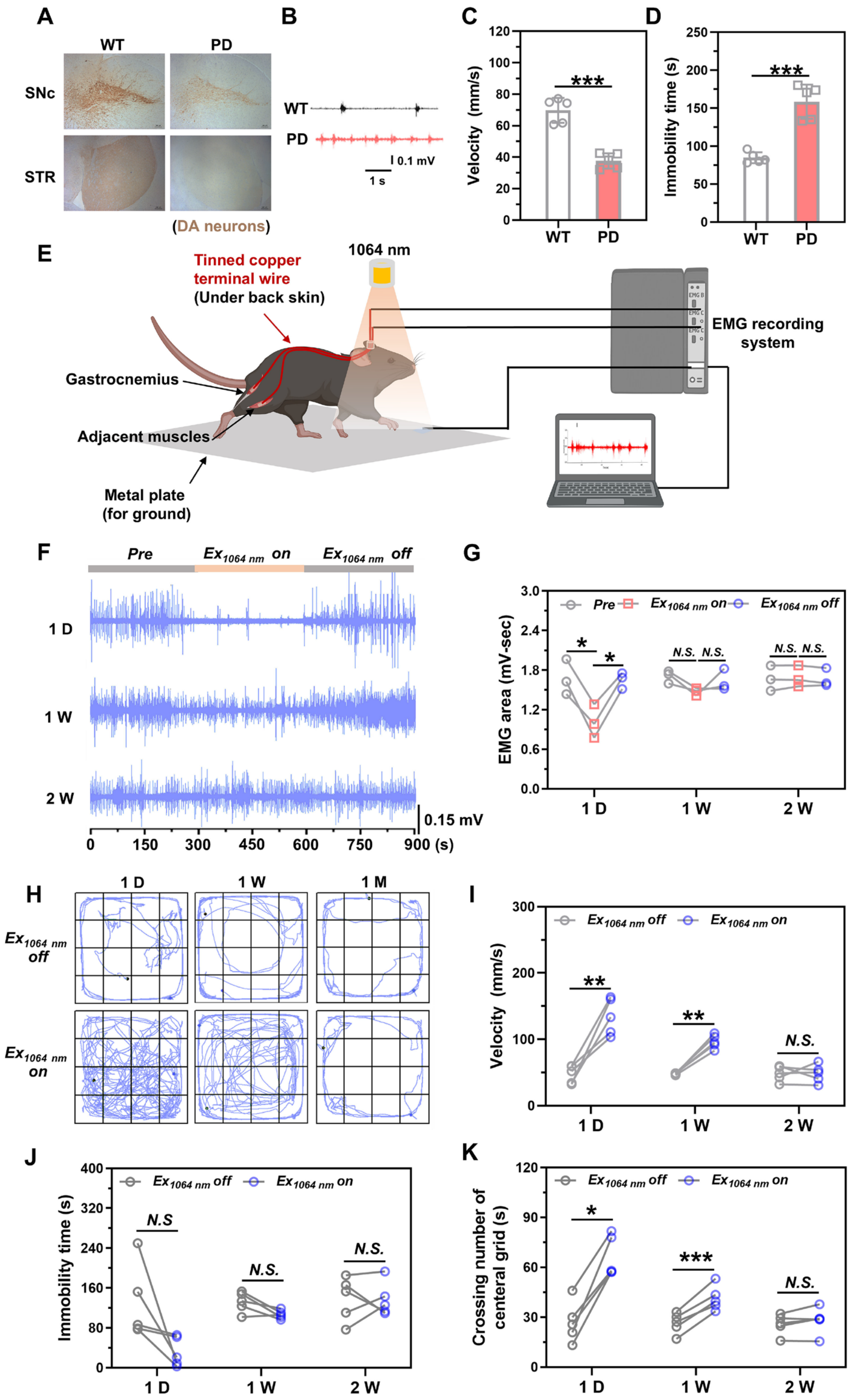


**Figure S7. Short-term and unstable intervention effects on tremor and motor dysfunction of PD mice treated by TRP-HPN under 1064 nm irradiation. (A-D)** Establishment and evaluation of PD mouse model: **(A)** IHC evaluation of DA neurons in SNc and STR regions of WT and PD mice. **(B)** EMG traces of WT and PD model mice. **(C-D)** Behavioral activity of PD mice, including velocity **(C)**, immobility time **(D)** (n=5). **(E)** Schematic diagram of EMG recording. Tinned copper terminal wire was fixed on the gastrocnemius, acting as recording electrodes of EMG, and the wires passed under the back skin and fixed to the head-plate without affecting mouse behaviors. **(F)** EMG traces from a PD mouse treated with TRP-HPN under 1064 nm irradiation (60 mW/cm$^2$, 5 min), recorded within 2 weeks post-delivery. **(G)** Statistical analysis of EMG area across groups (n=3). **(H)** Movement trajectories of PD TRP-HPN mice under 1064 nm irradiation. **(I and K)** Quantification of velocity **(I)**, immobility time **(J)**, and time spent crossing the center grids **(K)** of PD TRP-HPN mice under 1064 nm irradiation (n=5). *P < 0.05, **P < 0.01, ***P < 0.001. Statistical data were presented as Mean ± S.D.

**Table S1. Parameter comparison of current emerging non-invasive neuronal stimulation techniques *in vivo*.**

| Stimulation technique *in vivo* | Power of irradiation for *in vivo* neuromodulation | Neurostimulation-responsive module | Wireless for *in vivo* study | Depth | Neuron-targeted activation | Latency time of behavioral modulation | Time scale | Reference |
|---|---|---|---|---|---|---|---|---|
| Optical stimulation | ***60 m W/cm²*** ***(Continuous 1064 nm)*** | ***HaloNeu (NIR-II sensitive hybrid channels)*** | ***Wireless*** | ***10 mm (60 mW/cm²)*** | ***Target*** | ***6.0 s*** | ***2 months*** | ***This work*** |
| | 1.5 $W/cm^2$ (808 nm, 20 Hz) | ATB NPs (Photothermal) | Wireless | 4.3 mm | Antibody targeting | N/A | N/A | *Sci. Adv.*, **11**, eado4927 (2025). |
| | 0.8-1 $W/cm^2$ (continuous 1064 nm) | MINIS (Photothermal) | Wireless | 4.5 mm | N/A | 5.0 s (0.5 mm depth) | N/A | *Nat. Biomed. Eng.*, **6**, 754-770 (2022). |
| | 161 $mW/cm^2$ (980 nm at 200 ms) | God nanorods (Photothermal) | Wireless | Retinas | Antibody targeting | N/A | Retinas for restoring light sensitivity | *Science*, **368**, 1108-11103 (2020). |
| | 80 $W/cm^2$ (635 nm, 20 Hz) | Optogenetics (ChRmine) | Wireless | 7 mm | Target | N/A | N/A | *Nat. Biotechnol*, **39**, 161-164 (2021). |
| | 1 $W/cm^2$ (653 nm, 1 Hz) | Drop-printed SiHF (Optoelectrical) | Wireless | Brain surface | N/A | N/A | N/A | *Science*, **389**, 1127-1132 (2025). |
| | 39.2 $W/cm^2$ (980 nm, 20 Hz) | HUP (Optoelectrical) | Wireless | 4.2 mm | N/A | N/A | 7 days | *Sci. Adv.*, **11**, eadt4771 (2025). |
| | 140 $W/cm^2$ (980 nm, 15 ms pulses at 20Hz) | UCNPs (Opsin activation) | Wireless | 0.5 mm | Target | N/A | N/A | *Science*, **359**, 679-684 (2018). |
| Magnetic stimulation | 10 mT, 100 Hz AMF | MENDs (Magnetoelectric) | Wireless | 4.5 mm | N/A | N/A | 2 weeks | *Nat. Nanotechnol.*, **20**, 121-131 (2025). |
| | ≥20 mT, ΔB<10T/m AMF | m-Torquer (Magnetomechanic) | Wireless | 4.75 mm | Antibody targeting | N/A | 2 weeks | *Nat. Nanotechnol.*, **19**, 1333-1343 (2024). |
| | 80 mT, 49.9 kHz AMF | SPIONs (Magnetothermal) | Wireless | *Drosophila* | N/A | 0.5 s | Acute application | *Nat. Mater.*, **21,** 951-958 (2022). |
| Acoustic stimulation | 1.40 MPa, 1.5 MHz | HOFs (Sono-chemogenetics) | FUS | 4.5 mm | Target | 4.0 s | 5 days | *Nature*, **638**, 401-410 (2025). |
| | 2.8 MPa, 0.6 MHz | Holographic transcranial ultrasound | Multi-element ultrasound | Sensory cortex | N/A | N/A | Acute application | *Nat. Biomed. Eng.*, (2025). https://doi.org/10.1038/s41551-025-01449-x. |
| | 1.92 MPa, 650 kHz | MiniUlTra | tFUS | 10 mm | N/A | N/A | 28 days | *Nat. Commun.*, **16**, 4940 (2025). |
| | 100 kPa, 50 kHz | ImPULS | Chronic implanted | 4.4 mm | N/A | N/A | N/A | *Nat. Commun.*, **15**, 4601 (2024). |
| Other stimulation | 613 nm (27.55 $W/cm^2$, 10 Hz) | Transcranial brain modulator LED | Immobilized on mouse | Multi-region stimulation | N/A | ms | N/A | *Nat. Commun.*, **15**, 10423 (2024). |
| | Microwave 1.7 $W/cm^2$, 2.05 GHz | A microwave split-ring resonator (Heating) | 2-4 mm (Outer diameter) | Cortex surface | N/A | N/A | N/A | *Sci. Adv.*, **10**, eado5560 (2024). |
| | N/A | Organic semiconducting oligomers | Wireless | *Hydra vulgaris* | N/A | N/A | N/A | *Sci. Adv.*, **9**, eadi5488 (2023). |

**Abbreviation:** ATB NPs: Au@TRPV1@β-syn nanoparticles; MINIS: Macromolecular infrared nanotransducers for deep-brain stimulation; Si-HF: Si heterojunction films;

HUP: Hybrid upconversion and photovoltaic nanoparticles; PIN-Si: p-i-n type Si; UCNPs: Lanthanide-doped upconversion nanoparticles; MENDs: Magnetoelectric nanodiscs; AFM: Alternating magnetic field; SPIONs: Superparamagnetic iron oxide nanoparticles; HOFs: Hydrogen-bonded organic frameworks; FUS: Focused ultrasound; MiniUlTra: Miniaturized and Bioadhesive coupled ultrasound transducers; tFUS: Transcranial focused ultrasound; ImPULS: Implantable piezoelectric ultrasound stimulator.
